# Mammalian Cells Integrate Endoplasmic Reticulum and Nuclear Envelope signals to time mitotic entry

**DOI:** 10.64898/2025.12.12.693814

**Authors:** Yusuke Takagi-Shiozaki, Nicholas Codallos, Natasha Saik, David Jenkins, Pablo Lara-Gonzalez, Andrew K. Shiau, Katharine S Ullman, Maho Niwa

## Abstract

Accurate cell division requires coordination between organelle organization and cell-cycle progression, but how architectural and functional cues from the endoplasmic reticulum (ER) and nuclear envelope (NE)—a continuous membrane network—interface with mitotic control remains unclear. Here, we demonstrate that mammalian cells integrate ER/NE structure and functions to regulate the onset and progression of mitosis. Perturbing ER function with diverse stressors causes a selective delay at the metaphase–anaphase transition, accompanied by defective spindle assembly, chromosome misalignment, and loss of coordinated ER–chromosome organization. Under these conditions, the checkpoint protein MAD1 fails to efficiently dissociate from the NE. ER stress also disrupts microtubule-organizing centers and the *centriculum*, an ER-derived compartment surrounding centrosomes. Restoring ER structure by expressing the shaping proteins CLIMP63(1–192) or REEP4 rescues spindle organization and mitotic progression. Conversely, transient metaphase arrest induced by partial APC/C inhibition remodels ER morphology independently of stress, and this remodeling is reversed by CLIMP63(1–192). These findings uncover a bidirectional link between ER structure function and the spindle assembly checkpoint, identifying the organelle architecture as an instructive signal that modulates mitotic timing in mammalian cell.

## Introduction

The endoplasmic reticulum (ER)----the entry point to the secretory pathway--- is a vast and dynamic organelle, lies at the center of eukaryotic cell organization (Schwarz and Blower, 2016). Structurally, the ER forms an extensive, continuous network of sheets and tubules that is physically connected to the nuclear envelope (NE), a membrane system consisting of an inner nuclear membrane (INM) and outer nuclear membrane (ONM), that are bridged by nuclear pore proteins complex (NPC). Specifically, in mammalian cells, the ER extends outward to form an elaborate reticular network that weaves throughout the cytoplasm (Matlack et al., 1998; Meldolesi and Pozzan, 1998; Puhka et al., 2007; Voeltz et al., 2002; Westrate et al., 2015; Zheng et al., 2022). Beyond its remarkable architecture and its unique position, the ER serves as the command center for the secretory pathway, directing the synthesis, folding, modification, and maturation of nearly all secreted and membrane-associated proteins(English and Voeltz, 2013; Schwarz and Blower, 2016; Wenzel et al., 2022). It also initiates the biosynthesis of lipids, laying the foundation for membrane production and cellular compartmentalization (Celik et al., 2023; Jacquemyn et al., 2017; Kim and Burd, 2023; Kwiatek et al., 2020; Radanovic and Ernst, 2021; Stevenson et al., 2016). As the functional demands of the ER fluctuate constantly during the life time of the cell in response to developmental and environmental cues, which are, in part, recognized and regulated by the Unfolded Protein Response (UPR) pathway. In *Saccharomyces cerevisiae*, the increased demands of ER homeostasis (collectively called ER stress) is recognized by IRE1, an UPR initiating component, which triggers transcriptional programs ultimately up-regulating chaperone expression, folding capacity, and lipid biosynthesis, thereby meeting the increased demands of the ER homeostasis. In metazoans, the UPR is mediated by three sensors—IRE1, ATF6, and PERK—which collectively increase ER functional capacities and lipid synthesis upon increasing components that increase protein folding capacities (Hetz et al., 2020; Hwang and Qi, 2018; Walter and Ron, 2011). Ultimately, cells deficient in UPR capacities undergo cell death, underscoring the importance of the signaling pathways like UPR. Furthermore, the importance of ER functional homeostasis is illustrated by human diseases caused by its mis-regulation such as diabetes, neurodegenerative diseases, and cancer (Jiang et al., 2016; Yuan et al., 2024; Zhang et al., 2024).

In addition to its functional significance, the ER cannot be generated ***de novo***, suggesting the presence of a mechanism for ensuring that cells will receive some levels of the ER during the cell cycle. In yeast, *S. cerevisiae*, we identified a cell cycle checkpoint which we termed the ER stress surveillance (ERSU) pathway, which is wired independently from the UPR pathway (Babour et al., 2010; Chao et al., 2019; Niwa, 2020; Pina et al., 2018; Pina et al., 2023; Pina and Niwa, 2015). We discovered that when ER is stressed or the ER is not sufficient for splitting into two cells during the cell division, the inheritance of the ER into the daughter cell is blocked temporarily, septin ring critical for cytokinesis is re-localized from the bud neck to the bud scar, a previous cell division site, ultimately leading to the cytokinesis block. These events are wired independently from the UPR, and thus we termed the ER Surveillance (ERSU) pathway. In addition to the ERSU pathway, ER stress also activates the UPR, allowing re-establishment of the ER functional homeostasis, at which point, the ER inheritance block is released and cells re-enter cell cycle. Furthermore, cells unable to halt cell division in response to ER stress, ultimately resulting in cell death, illustrating the significance of the ERSU cell cycle checkpoint, in addition to the UPR.

While UPR is conserved in mammalian cells, we do not yet know if cell cycle mechanisms, such as the ERSU pathway, exist to integrate functional capacities of the ER into mitosis. Furthermore, molecular mechanisms of mitosis in mammalian cells differ significantly from those in yeast. In mammalian cells, the ER forms an extensive, continuous network of sheets and tubules that is physically connected to the nuclear envelope (NE), a membrane system consisting of an inner nuclear membrane (INM) and outer nuclear membrane (ONM) that are bridged by nuclear pores.

During mitosis, the NE breaks down (NEBD) and INM components merge with the ER and both lipids and INM proteins are distributed throughout the ER. In addition to certain nuclear pore complex components such as the Y-complex (Imamoto and Funakoshi, 2012; Zuccolo et al., 2007), NE dynamics also regulates certain key mitotic factors including MAD1, 2 (Cunha-Silva and Conde, 2020; Mossaid and Fahrenkrog, 2015), and CENP-F (Berto and Doye, 2018; Zuccolo et al., 2007), which are essential for establishing metaphase chromatin organization. Thus, after NEBD during mitosis, the ER becomes to directly face the condensing or the condensed chromatin, which in turn associates with centrosomes, kinetochores, and the spindle—a relationship essential for subsequent NE reassembly once chromosomes segregate (Guttinger et al., 2009; Kutay et al., 2021). This cyclical redistribution suggests that ER integrity could influence chromosome organization and NE formation. Indeed, during metaphase after NE disassembly, condensed metaphase chromosomes are positioned directly interfacing with surrounding ER membranes, without NE insulating the genome from the ER. While ER stress has been linked to G1/S phase arrest via PERK-mediated phosphorylation of eIF2α which diminishes translation initiation efficiencies and resulting in suppression of cyclin D1 translation (Brewer and Diehl, 2000; Cabrera et al., 2017; Niwa and Walter, 2000). As GADD34 phosphatase dephosphorylates eIF2α, cyclinD1 levels increase, resulting in the release from G1/S phase block and re-entry into the cell cycle. Interestingly, the functional impact of ER stress on later mitotic stages remains largely unexplored.

Here in our study, we dissected how ER homeostasis influences mitosis in mammalian cells by inducing ER stress during cell division. We found that ER stress delayed the metaphase-to-anaphase transition and disrupted the spatial organization of the ER around mitotic chromosomes. Overexpression of ER-structural proteins REEP4 or CLIMP63(1–192) restored normal timing, revealing a regulatory role for ER architecture in mitotic progression. ER stress also perturbed chromosome alignment, spindle organization, centrosome separation, nuclear envelope breakdown, and the release of the checkpoint protein MAD1 from the NE. Moreover, ER stress disorganized the integrity of the centriculum—an ER-derived membrane encircling the centrosome—suggesting a mechanistic link between ER structure and spindle biogenesis. Conversely, mitotic delay induced by inhibition of the anaphase promoting complex/cyclosome (APC/C) degradation by MG132, a proteasome inhibitor, resulted in disruption of the ER organization, suggesting that ER structure homeostasis is an integrated part of the indicating that ER structure and mitotic timing are mutually regulated processes, illustrating the active role of the ER in mitosis.

## Results

### ER stress delays the metaphase–anaphase transition

To investigate how endoplasmic reticulum (ER) homeostasis influences cell division, we examined how ER stress alters the dynamics of mitotic progression. Using U2OS cells stably expressing mEmerald–Sec61β, a fluorescent reporter for the ER translocon complex, we visualized ER morphology while simultaneously tracking chromatin with NucBlue or SiR-DNA (**Figure 1A**). ER stress was induced with tunicamycin (Tm), an inhibitor of N-linked glycosylation known to cause the accumulation of misfolded proteins in the ER. Time-lapse fluorescence imaging then allowed us to monitor, in real time, how disruption of ER homeostasis remodels ER architecture and impacts mitotic behavior.

**Figure 1.**
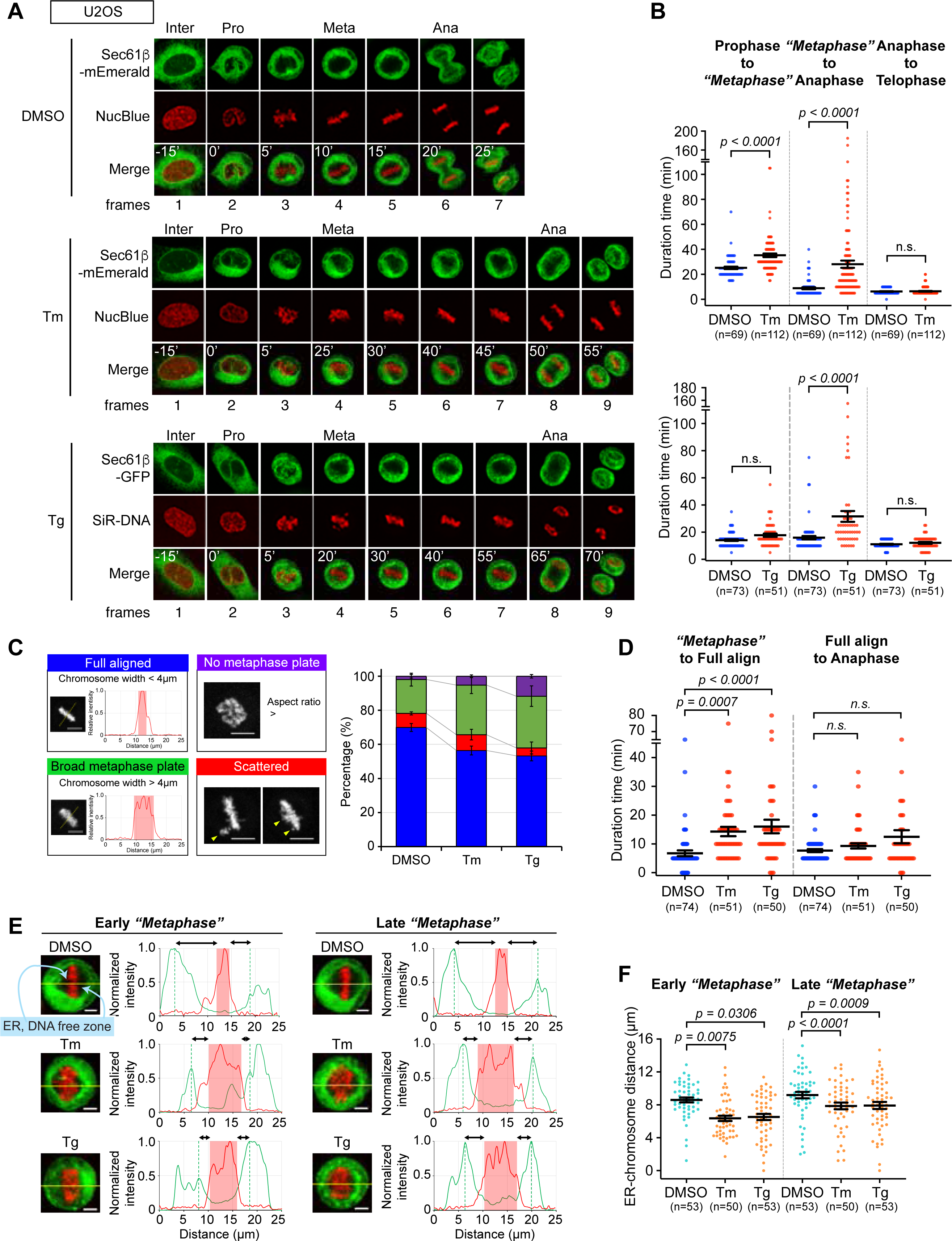
ER stress slows the metaphase–anaphase transition and disrupts ER organization around mitotic chromosomes. (**A**) To visualize ER dynamics during mitosis, U2OS cells stably expressing mEmerald–Sec61β were stained with NucBlue and treated with DMSO or a well-established ER stress inducer, tunicamycin (Tm; 10 μg/mL) for 6 h before spinning-disk confocal time-lapse imaging every 5 min for 16 h. In parallel, U2OS cells expressing GFP–Sec61β were stained with SiR-DNA, treated with DMSO or another ER stress inducer, thapsigargin (Tg; 200 nM) for 6 h, and imaged every 5 min for 10 h. Prophase onset, identified by chromosome morphology, was set as time 0 min. Scale bars, 10 μm. (**B**) Quantification of mitotic timing revealed that ER stress prolongs specific transitions within mitosis. Phase durations were measured from asynchronously dividing cells in three independent time-lapse experiments, analyzing all mitotic cells in each field (n). Data represent mean ± SEM from individual cells, and statistical significance was determined by one-way ANOVA with Sidak’s multiple comparisons test. (**C**) To examine chromosome congression defects, metaphase alignment was scored one frame before anaphase onset. Cells were grouped by alignment phenotype: fully aligned (blue), broad metaphase plate (green), incomplete plate (purple), or scattered plate (red). The frequency of each category is plotted, revealing increased misalignment under ER stress. (**D**) The timing between metaphase chromosome alignment and the onset of anaphase was further measured. Under ER stress, even cells forming a metaphase plate immediately prior to anaphase frequently exhibited broad plates (>4 μm width), indicating partial or unstable alignment. (**E**) ER organization relative to condensed chromosomes was analyzed by line-scan fluorescence profiles along the yellow line in “*metaphase”* (or prometaphase/metaphase) frames. Normalized intensity plots of mEmerald–Sec61β (ER, green) and NucBlue (chromosomes, red) illustrate the distance between the chromosome region (red-shaded) and the nearest ER fluorescence peak (>50% of maximum; green dashed lines). Scale bars, 10 μm. (**F**) Quantification of these ER–chromosome distances across all mitotic cells confirmed that ER stress increases the spatial separation between the ER and metaphase chromosomes, reflecting altered ER distribution during mitosis.

Mitotic stages and their durations were defined based on chromatin condensation dynamics and compared between unstressed (DMSO-treated) and ER-stressed (tunicamycin-treated) cells (**Figure 1A–B**). Prophase was marked by the onset of chromatin condensation, metaphase by chromosome alignment at the metaphase plate, and anaphase by the initiation of sister chromatid separation. Under normal conditions, most cells spent approximately 10–15 minutes in metaphase, with chromosomes tightly aligned within ∼4 µm prior to anaphase onset, followed by chromatin de-condensation as the nuclear envelope (NE) reassembled (**Figure 1C**). In contrast, ER-stressed cells showed a pronounced delay in the metaphase-to-anaphase transition, while the durations of earlier (prophase–metaphase) and later (anaphase–telophase) stages remained comparable to controls. These findings suggest that ER stress selectively perturbs mitotic timing at the metaphase–anaphase transition. Consistent with this, similar metaphase delays were observed when ER stress was induced by thapsigargin (Tg), an inhibitor of the ER Ca²⁺-ATPase SERCA (**Figure 1A–B**).

To determine whether this mitotic delay was cell type–specific, we analyzed HeLa cells stably expressing the ER reporter GFP–Sec61β and the chromatin marker mCherry–histone H2B. As observed in U2OS cells, tunicamycin (Tm) treatment in HeLa cells selectively prolonged the metaphase-to-anaphase transition, with minimal effect on other stages such as prophase-to-metaphase or anaphase-to-telophase (***Figure S1A–B***). A similar extension of metaphase duration occurred when ER stress was induced by β-mercaptoethanol (β-ME), which disrupts disulfide bond formation by altering ER redox balance. Together, these findings indicate that ER stress—regardless of the inducing agent—specifically lengthens mitotic progression by delaying the metaphase-to-anaphase transition, revealing a previously unrecognized link between ER homeostasis and chromosome segregation.

### Molecular mechanisms of metaphase-to-anaphase delay induced by ER stress

To understand how ER stress delays the metaphase-to-anaphase transition, we examined chromosomal organization during metaphase. In ER-stressed cells, metaphase chromosomes appeared broader and less compact than in controls (**Figure 1A, C**). Under DMSO-treated conditions, approximately 70.7% of cells achieved a tightly aligned metaphase plate, whereas only about 50% of tunicamycin- or thapsigargin-treated cells reached this state (**Figure 1C, blue**). Many ER-stressed cells failed to form a compact metaphase plate within 4 µm, maintaining a broader and less organized chromosome arrangement even immediately before anaphase onset, despite prolonged metaphase duration. In contrast, ER-stressed cells that successfully achieved full chromosome alignment progressed through anaphase with normal timing. These results suggest that ER stress extends metaphase primarily by impairing chromosome alignment, potentially through altered mitotic ER organization or its interactions with spindle and chromosomal structures.

To more precisely define which step within metaphase was affected, we subdivided the metaphase–anaphase transition into two intervals: from pre-metaphase/metaphase entry to complete chromosome alignment, and from full alignment to anaphase onset. ER stress induced by tunicamycin or thapsigargin selectively extended the chromosome alignment phase, whereas the duration from full alignment to chromatid separation remained comparable between control and ER-stressed cells (**Figure 1D**). These findings demonstrate that ER stress primarily compromises chromosome alignment rather than anaphase onset. Throughout this study, pre-metaphase or metaphase cells in which chromosomes remained partially aligned—spanning a metaphase plate width greater than 4 µm—are collectively referred to as “*metaphase”* under ER stress to reflect their prolonged occupancy of these stages due to delayed spindle organization.

### Disruption of ER homeostasis led to striking changes in metaphase ER–chromosome organization

To determine whether additional alterations in pro-metaphase or metaphase architecture contributed to the extended “*metaphase*” duration, we examined how ER stress reshapes ER morphology and its spatial relationship with chromosomes. One hallmark of mammalian mitosis is nuclear envelope breakdown (NEBD), which enables spindle microtubules to access condensed chromosomes and bind kinetochores for faithful segregation (Espeut et al., 2015). Concurrently, the ER network—composed of tubules, sheets, and cisternae—undergoes characteristic mitotic remodeling, yielding a tubule-rich peripheral ER and a sheet- and cisternae-enriched perinuclear ER (Obara et al., 2023). Although the extent of this reorganization varies by cell type (Anderson and Hetzer, 2008; Lu et al., 2009; Merta et al., 2021; Puhka et al., 2007; Schlaitz et al., 2013; Zhao et al., 2023), DMSO-treated control cells consistently exhibited a distinct metaphase ER morphology in which ER membranes were excluded from the chromosome region, forming a well-defined ER-free zone around the metaphase plate (**Figure 1E**).

Upon ER stress induction, the organization and spatial relationships between the ER and metaphase chromosomes were markedly altered (**Figure 1E**). Because ER stress prolonged the metaphase-to-anaphase transition (**Figure 1B**), we focused on ER morphology and its positional relationship with condensed metaphase DNA at both early and late metaphase phases. Early metaphase was defined by the initial frame following chromosome alignment. Using line-scan analysis across the metaphase DNA axis, we quantified the relative fluorescence intensity profiles of ER and DNA signals. In control cells, early metaphase exhibited distinct, well-separated fluorescence peaks for ER and DNA (**Figure 1E–F, DMSO**). Similar spatial separation was observed later in metaphase, just prior to anaphase onset. In contrast, tunicamycin (Tm) treatment broadened the ER peaks and reduced the distance between ER and DNA compared with controls. As cells progressed to late metaphase, ER distributions flanking the metaphase DNA became more separated in both Tm- and thapsigargin (Tg)-treated populations; however, in stressed cells, ER remained consistently closer to the DNA than in unstressed U2OS cells (**Figure 1E–F**). We observed similar ER-metaphase chromatin and HeLa cells (***Figure S1C-D***). These results indicate that ER stress alters the spatial organization of the mitotic ER, positioning ER membranes abnormally close to condensed chromosomes throughout metaphase. This sustained proximity persisted through the extended metaphase interval in ER-stressed cells, contrasting with the greater ER–chromosome separation characteristic of unstressed conditions, and likely reflects a stress-induced remodeling of ER architecture during mitosis.

To further assess ER–chromatin interactions, we examined ER association with the peripheral regions of metaphase chromosomes. Binary masks were generated from segmented ER and DNA signals, and overlap was quantified specifically at the terminal 20% segments of the metaphase DNA (***Figure S1E***). Even under control conditions, U2OS cells exhibited limited overlap between ER and chromosome ends. However, ER-stressed cells displayed a pronounced increase in ER coverage at DNA termini, with significantly greater ER–DNA overlap throughout metaphase compared to controls (***Figure S1F–S1G***). A comparable pattern was observed in HeLa cells at both early and late metaphase (***Figure S1H–S1I***).

Collectively, these findings reveal that metaphase progression is accompanied by dynamic ER reorganization that is highly sensitive to ER homeostasis. Disruption of ER function or induction of ER stress leads to aberrant ER–chromosome positioning and sustained ER proximity to mitotic DNA, thereby perturbing proper chromosome alignment and delaying anaphase entry. These data underscore the active role of the ER in shaping mitotic chromosome architecture and maintaining coordinated organelle and genome organization.

### ER structural alterations restore ER stress–induced mitotic delay

Our findings thus far reveal that disruption of ER homeostasis profoundly disturbs spindle alignment, thereby delaying the critical metaphase-to-anaphase transition. In mammalian cells, the ER typically reorganizes from a sheet-like network during interphase into a highly branched tubular structure during mitosis (Puhka et al., 2012; Puhka et al., 2007; Voeltz et al., 2002; Wang et al., 2013). This transformation contrasts with the pattern observed in *Drosophila* and *C. elegans* embryos, where mitosis instead promotes an expansion of ER sheets (Puhka et al., 2007, 2012). Although such dynamic changes in ER architecture have been well documented, their molecular significance during mitosis remains largely unresolved (Bergman et al., 2015; Zhao et al., 2023). Based on our results, we hypothesized that ER homeostasis is integrated into mitotic control, influencing spindle assembly and chromosome segregation. Specifically, perturbing the transition of ER structure from interphase to metaphase, or altering the spatial relationship between ER membranes and sister chromatids, could determine the timing of anaphase onset. To test this model, we examined whether modulating ER morphology through overexpression of structural proteins such as CLIMP63 or REEP4 could influence mitotic timing.

We first assessed the effect of CLIMP63 overexpression on ER remodeling during mitosis. CLIMP63 (Cytoskeleton-Linking Membrane Protein, 63 kDa) promotes ER sheet formation by dimerizing through its luminal C-terminal domain, thereby converting tubules into sheets (Shibata et al., 2010) (**Figure 2A**). The C-terminal region contains multiple coiled-coil motifs—predicted to number either three (Klopfenstein et al., 2001; Sandoz and van der Goot, 2015; Shen et al., 2019) or five (Xu et al., 2023)—that regulate ER sheet width and lumen size. In contrast, the truncation mutant CLIMP63(1–192), lacking these motifs, acts dominantly to disrupt normal sheet morphology, producing narrowed lumens with enhanced curvature (Shen et al., 2019; Wang et al., 2022). Using time-lapse imaging, we found that overexpression of CLIMP63(1–192) markedly reduced the metaphase-to-anaphase delay induced by ER stress (**Figure 2C, lanes 2 vs 4**). Importantly, this truncation variant had no effect on timing in unstressed (DMSO-treated) cells (**Figure 2C, lanes 1 vs 3**), indicating that ER structural remodeling specifically modulates mitotic progression under ER stress conditions.

**Figure 2.**
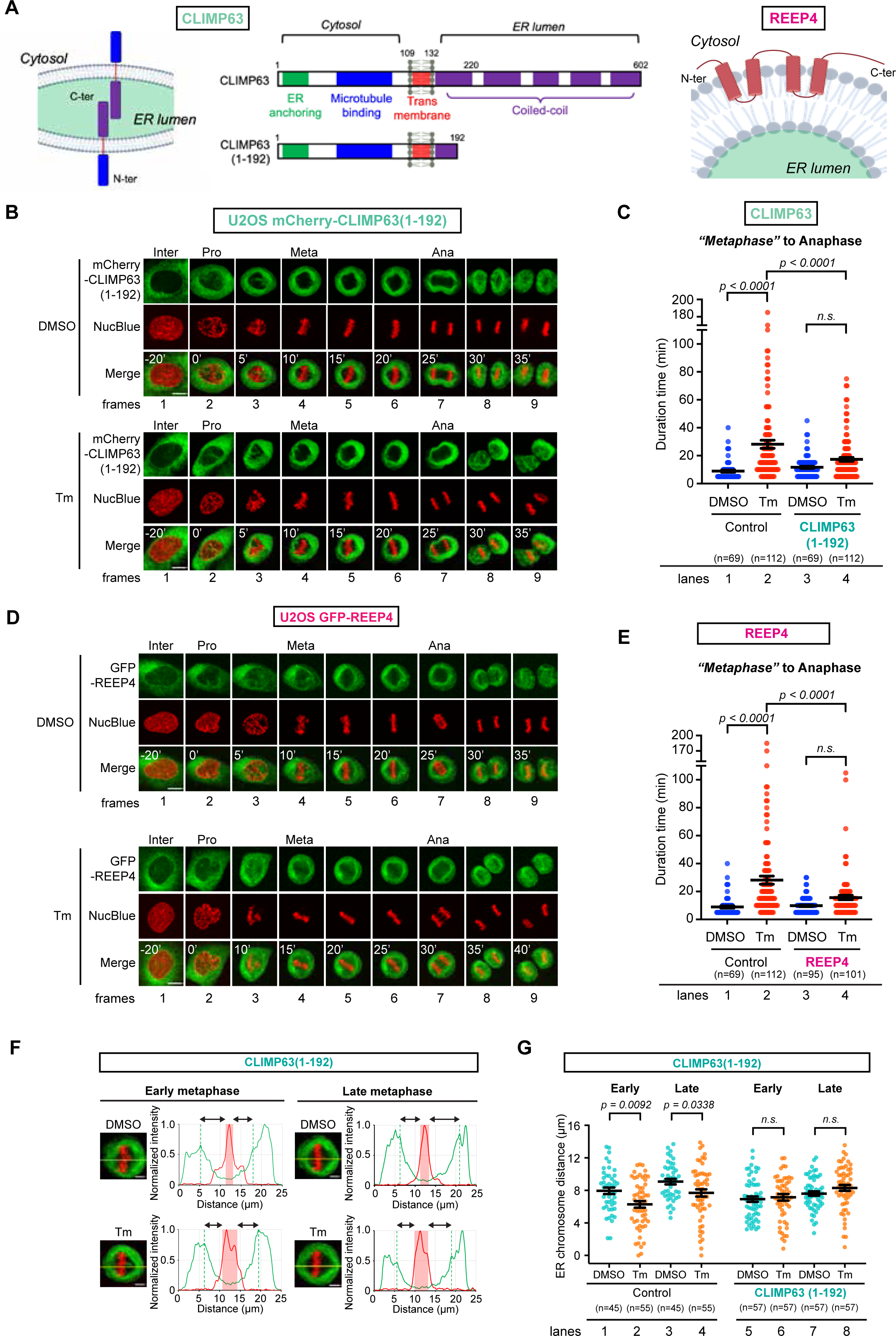
ER membrane–shaping proteins CLIMP63(1–192) and REEP4 alleviate ER stress–induced mitotic defects. (**A**) Schematic of CLIMP63 and REEP4 topology within the ER membrane, highlighting the luminal, transmembrane, and cytosolic domains. Full-length CLIMP63 and the N-terminal truncation CLIMP63(1–192) are shown to emphasize the region retained in the mitosis-protective construct. (**B**) Representative time-lapse sequences showing mitotic progression in U2OS cells stably expressing mCherry–CLIMP63(1–192) under control conditions or ER stress. Cells were stained with NucBlue, treated with DMSO or Tm (10 μg/mL), and imaged every 5 min for 16 h to visualize how CLIMP63(1–192) expression remodels ER organization during mitosis. Scale bars, 10 μm. (**C**) Metaphase-to-anaphase duration was quantified for the conditions shown in (B), using U2OS cells lacking CLIMP63(1–192) from Figure 1A as a non-expressing control. Data represent mean ± SEM from individual cells, and P values were determined by one-way ANOVA followed by Sidak’s multiple comparisons test; the same statistical approach was applied throughout the figure. (**D**) Representative time-lapse images of mitotic progression in U2OS cells stably expressing GFP–REEP4 under control or ER stress conditions. Cells were stained with NucBlue, treated with DMSO or Tm (10 μg/mL), and imaged every 5 min for 10 h, with prophase onset defined by the appearance of condensed DNA and set as time 0 min. Scale bars, 10 μm. (**E**) Metaphase-to-anaphase duration was measured for REEP4-expressing cells under the conditions in (D) and compared to cells lacking REEP4, revealing how REEP4 expression influences mitotic timing under ER stress. (**F**) Line-scan analysis of ER–chromosome organization at early and late metaphase in cells expressing CLIMP63(1–192), with or without Tm treatment. Normalized fluorescence intensities of mCherry–CLIMP63(1–192) (ER, green) and NucBlue (chromosomes, red) are plotted to illustrate how CLIMP63(1–192) expression reshapes ER distribution around metaphase chromosomes. Scale bars, 5 μm. (**G**) Quantitative analysis of ER signal overlap at metaphase chromosome ends in cells with or without CLIMP63(1–192) expression treated with DMSO or Tm. These measurements assess the extent to which CLIMP63(1–192) restores ER–chromosome contacts under ER stress. Scale bars, 5 μm.

To further define the spatial interplay between the ER and metaphase chromosomes, we performed line-plot analyses of ER and DNA distributions at early and late metaphase. In cells expressing CLIMP63(1–192), ER localization remained strikingly consistent across metaphase stages, in contrast to the dynamic redistribution observed in control cells (**Figure 2F–G**). Under ER stress, CLIMP63(1–192) expression also reduced ER accumulation at the distal ends of metaphase chromosomes (***Figure S2A–B***). In Tm-treated controls, ER overlap at chromosome termini was markedly elevated, whereas this enrichment was significantly attenuated by CLIMP63(1–192) overexpression (***Figure S2B–C,*** lanes 1–2 and 3–4 vs 5–8). Notably, the reduction in ER–chromosome overlap correlated with a shortened metaphase-to-anaphase transition, underscoring the functional link between ER architectural integrity and mitotic progression (**Figure 2C**, lanes 1–2 vs 3–4).

### REEP4 overexpression mitigates ER stress effects on mitosis

To further define how ER architecture governs mitotic progression, we examined another class of ER-shaping proteins by overexpressing REEP4, a reticulon family member containing a conserved reticulon homology domain (RHD). REEP3 and REEP4 were previously shown to maintain a critical exclusion zone between mitotic ER and metaphase chromatin, though the underlying molecular mechanisms remain elusive (Golchoubian et al., 2022; Kumar et al., 2019; Schlaitz et al., 2013). Motivated by the ER–chromosome repositioning observed under ER stress (**Figure 2D–E**), we performed time-lapse imaging of U2OS cells stably expressing REEP4 under the same perturbation conditions used for CLIMP63(1–192).

REEP4 overexpression did not alter the metaphase-to-anaphase interval in unstressed cells (**Figure 2D–E**, lanes 1 vs 3), consistent with normal mitotic timing. In contrast, under ER stress, REEP4 expression shortened the prolonged metaphase duration and restored inter-chromosomal ER spacing to levels comparable with DMSO-treated controls (**Figure 2E**, lanes 1–2 vs 3–4). Likewise, the ER stress–induced compression of ER–DNA spacing at both early and late metaphase (***Figure S2C–D***, lanes 1–2 and 3–4) was reversed by REEP4 overexpression (lanes 1–2 vs 5–6 for early metaphase; lanes 3–4 vs 7–8 for late metaphase). These effects closely paralleled the rescue observed with CLIMP63(1–192) (**Figure 2F–G**). Together, these findings identify ER morphogens such as CLIMP63(1–192) and REEP4 as key stabilizers of mitotic fidelity under stress, indicating that targeted remodeling of ER structure can preserve both metaphase timing and chromosome segregation in the presence of ER stress.

### ER stress impairs mitotic timing and alignment through spindle disruption

Overexpression of ER-structural regulators such as CLIMP63(1–192) or REEP4 rescued the ER stress–induced extension of the metaphase-to-anaphase interval (**Figure 2**), primarily by reducing the proportion of cells with abnormally broad metaphase plates (**Figure 3A**; decrease in green, lanes 2 vs 4 or 6, and corresponding increase in blue). These results suggest that delayed mitosis under ER stress arises from impaired chromosome alignment, and that preserving ER structural homeostasis can partially re-establish proper metaphase chromatin positioning, thereby restoring timely anaphase onset.

**Figure 3.**
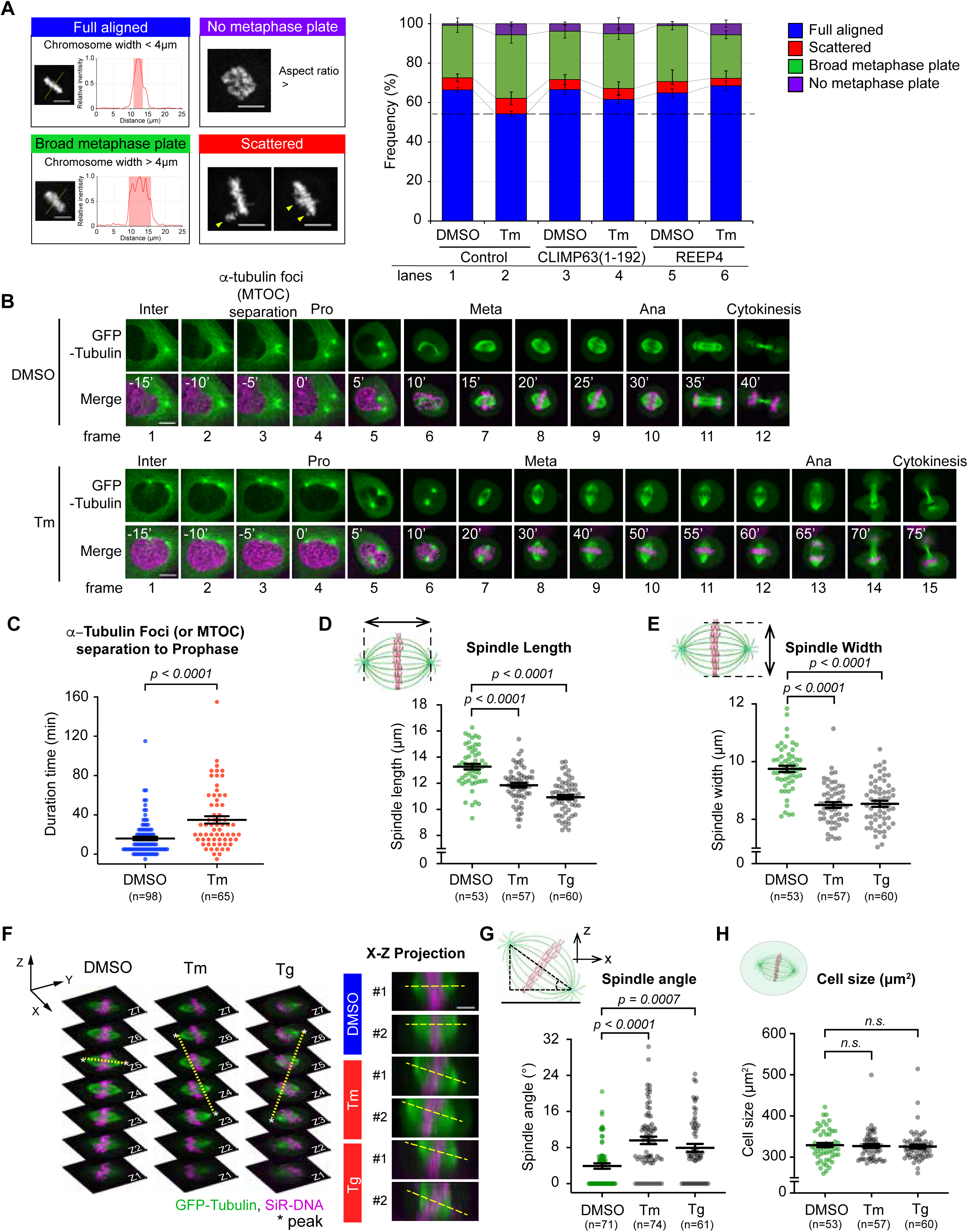
ER stress disrupts spindle assembly and metaphase architecture. (**A**) Metaphase chromosome organization, scored one frame before anaphase in ER-stressed U2OS cells, was categorized as in Figure 1C. Cells with fully aligned chromosomes and a metaphase plate <4 µm are shown in blue, cells with a broad metaphase or “metaphase” plate in green, cells lacking a clear plate or displaying only a broad plate in purple, and cells with scattered chromosomes in red; DMSO- or Tm-treated cells over-expressing CLIMP63(1–192) (lanes 3, 4) or REEP4 (lanes 5, 6) were compared with non-expressing controls (lanes 1, 2). (**B**) Representative time-lapse sequences show mitotic progression in U2OS cells stably expressing GFP–α-tubulin under control or ER stress conditions. Cells were stained with SiR–DNA, treated with DMSO or Tm for 6 h, and imaged every 5 min for 10 h, with prophase onset defined as time 0 min. Scale bars, 10 µm. (**C**) The interval from division of the GFP–α-tubulin focus/microtubule-organizing center (MTOC) to prophase onset was quantified. The MTOC was defined as a bright GFP–tubulin focus that resolved into two distinct foci at the time of division. (**D, E, G, H**) Metaphase spindle architecture was analyzed in U2OS cells by measuring spindle length (pole-to-pole distance, **D**), spindle width (maximum span of microtubule bundles, **E**), spindle angle (**G**), and cell size (**H**). Metaphase images were taken one frame before anaphase from time-lapse series acquired under the conditions in (**B**), including parallel experiments with Tg (200 nM); scale bars, 5 µm. P values were calculated using one-way ANOVA followed by Sidak’s multiple comparisons test, and in (A) means were derived from three independent experiments; data are presented as mean ± SEM from individual cells. (**F**) Z–X views of GFP–α-tubulin and metaphase chromosomes in U2OS cells treated with DMSO, Tm, or Tg illustrate alterations in spindle orientation. Spindle angles in the Z–X plane were quantified for each condition (right), with yellow dotted lines marking the connection between the two GFP–α-tubulin/MTOC peaks.

To determine how ER stress compromises chromosome alignment, we visualized microtubule dynamics using a stable GFP–α-tubulin–expressing U2OS line. Time-lapse imaging tracked microtubule-organizing centers (MTOCs)/centrosome behavior during mitosis (**Figure 3B–C**). Since α-tubulin is nucleated at centrosomes or microtubule-organizing centers (MTOCs) and can be visualized as distinct foci during the G2 phase, the spatial and temporal behavior of α-tubulin foci provides a proxy for centrosome-related information such as spatial positioning and separation.

In control cells, duplicated α-tubulin foci appeared during prophase and gradually separated along opposite poles as mitosis progressed (**Figure 3B**, frame 3, DMSO). Strikingly, in ER stressed cells, MTOCs were often visible earlier, even during interphase, and separated prematurely (**Figure 3B**, frame 1, Tm-treated). Similar phenotypes have been reported in yeast, where ER stress accelerates duplication of the spindle pole body (SPB)—the functional homolog of the yeast MTOC or centrosome (Lai et al., under review). These observations suggest that ER stress perturbs spindle pole duplication and positioning in a conserved manner across species.

Next, we quantified the timing between initial MTOC separation and prophase onset. In unstressed cells, α-tubulin foci separation occurred approximately 15 minutes before prophase, whereas in Tm-treated cells, this interval extended to ≥ 35 minutes, with more than 20% of cells showing delays exceeding 60 minutes (**Figure 3C, *S3A***). This prolonged interval indicates disrupted centrosome dynamics under ER stress conditions.

Proper spindle formation requires precise MTOC positioning during mitosis. In control cells, elongated α-tubulin fibers formed symmetric spindle structures aligned with metaphase chromosomes (**Figure 3B**, frame 10, DMSO). ER-stressed cells, however, exhibited distorted spindles (**Figure 3B**, frame 13, Tm-treated). To quantify this, we measured spindle pole-to-pole distance (spindle length) and the width of the α-tubulin bundles (spindle width) at one frame before anaphase onset (**Figure 3D–E, *S3B–C***). Both parameters were significantly reduced under ER stress. Additionally, we noticed that MTOCs and metaphase chromatin were coplanar in unstressed cells but misaligned across z-planes in stressed cells (**Figure 3F**, two tubulin peaks were positioned at z5-z5 in DMSO-treated cells, but shifted to z6-z3 in Tm or Tg treated cells). Measuring the angle between chromatin and the MTOC connection line (“spindle angle”) confirmed a significant increase in spindle tilt under ER stress (**Figure 3G** and ***S3C***).

Because spindle size scales with cell size (Rieckhoff et al., 2019), we assessed whether reduced spindle dimensions reflected smaller cell size. Using GFP background signal from α-tubulin to define cell boundaries, as described in previous studies (Kletter et al., 2022) (Yamagata and FitzHarris, 2013), we found that cell area was comparable between unstressed and ER-stressed cells (**Figure 3H** and ***S3D***). Thus, the spindle defects observed under ER stress reflect intrinsic perturbations of mitotic architecture rather than cell size differences.

### ER homeostasis maintains centrosomal architecture

Live-cell imaging revealed that tubulin foci duplicated at unexpectedly rapid rates, suggesting a regulatory interplay between centrosomes and the ER. This observation prompted a closer examination of spatial relationships reminiscent of those described in *C. elegans* embryos, where the centrosome docks along the nuclear envelope (NE) in tight association with ER membranes shortly after fertilization (Araujo et al., 2023; Luders, 2023; Maheshwari et al., 2023; Woodruff, 2021).

This intimate ER–centrosome relationship is essential for maintaining spatial fidelity: mutations that disrupt their alignment consistently cause abnormal centrosome duplication and mispositioning during G2, ultimately predisposing cells to mitotic defects.

Guided by this framework, we visualized MTOCs and ER organization under ER stress (**Figure 4**). In control cells, DMSO treatment preserved a well-defined structure in which the ER formed a sharp, continuous rim encircling each α-tubulin focus (**Figure 4A**). Upon Tm treatment, however, this organization deteriorated—ER membranes became diffuse and irregular, losing the distinct periphery marking the MTOC (**Figure 4B**). These changes indicate that ER stress disrupts the precise ER–MTOC interface, potentially destabilizing early centrosome positioning and duplication. Strikingly, such alterations appeared before mitotic onset, suggesting that ER stress initiates a cascade culminating in spindle defects, chromosome misalignment, and extended metaphase duration. This sequence suggests a mechanistic link between ER structural homeostasis and mitotic fidelity.

**Figure 4.**
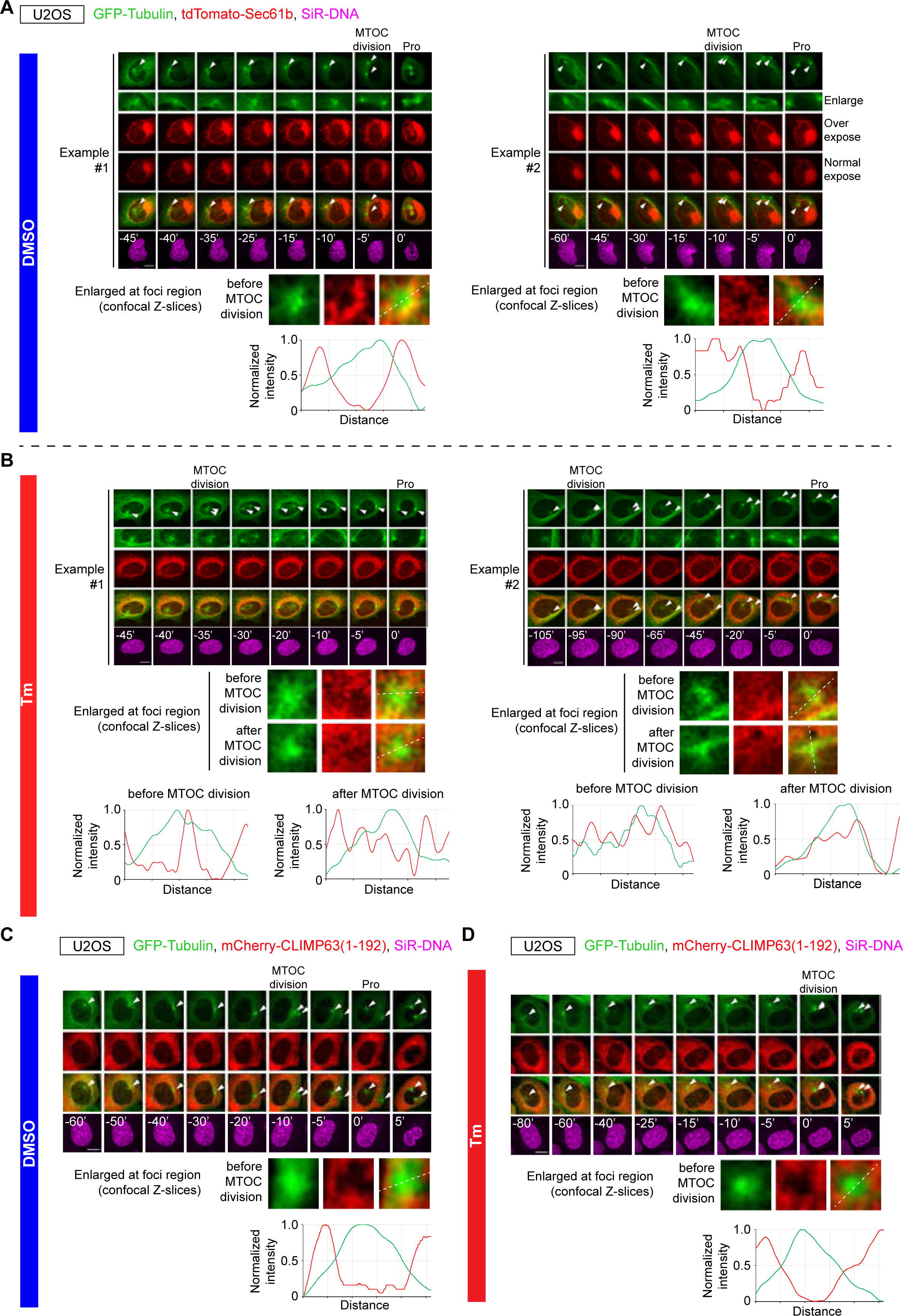
ER stress perturbs ER organization around microtubule-organizing centers (MTOCs). (**A, B**) Time-lapse imaging of ER distribution around MTOCs in U2OS cells under control conditions (DMSO, A) or ER stress (Tm, B). Cells stably expressing GFP–α-tubulin and tdTomato–Sec61β were stained with SiR–DNA, treated with DMSO or Tm (10 µg/mL) for 6 h, and imaged over time, with prophase onset defined as time 0 min. Arrowheads mark MTOCs, and enlarged views highlight ER organization before and after MTOC division, accompanied by line-scan fluorescence intensity profiles of ER (red) around MTOCs. Scale bars, 10 µm. (**C, D**) Effect of CLIMP63(1–192) overexpression on ER organization around MTOCs in the presence or absence of ER stress. U2OS cells co-expressing GFP–α-tubulin and mCherry–CLIMP63(1–192) were treated with DMSO (C) or Tm (D) and imaged by time-lapse microscopy to visualize how CLIMP63(1–192) modulates ER distribution surrounding MTOCs under both unstressed and Tm-treated conditions. Scale bars, 10 µm.

To test whether these phenotypes are intrinsic to ER organization, we examined whether CLIMP63(1–192) overexpression could restore ER–MTOC alignment under stress. In unstressed cells, CLIMP63(1–192) expression did not alter ER–MTOC spatial organization, which remained comparable to control architecture (**Figure 4C**). Under ER stress, however, CLIMP63(1–192) expression strikingly reinstated ER integrity around the α-tubulin foci, reestablishing the structured network characteristic of the *centriculum*—the organized interface between ER membranes and spindle microtubule foci—confirmed by line-plot analysis (**Figure 4D**). These results demonstrate that maintaining ER structural integrity is sufficient to preserve ER–MTOC positioning during mitosis, underscoring ER homeostasis as a central determinant of spindle organization and mitotic accuracy.

### ER stress delays NEBD and MAD1 release

Based on the spindle assembly phenotype, we next investigated molecular events that govern microtubule attachment to sister chromatids in addition to effects on the MTOC. Because the ONM is continuous with the ER, ER stress could perturb NE dynamics. To test this, we performed live-cell imaging of U2OS cells stably expressing the NPC protein POM121 fused to mNeonGreen (POM121–mNG), together with SiR-DNA to label chromatin, and monitored NE behavior in unstressed and ER-stressed cells (**Figure 5A–C**, and ***S4A***).

**Figure 5.**
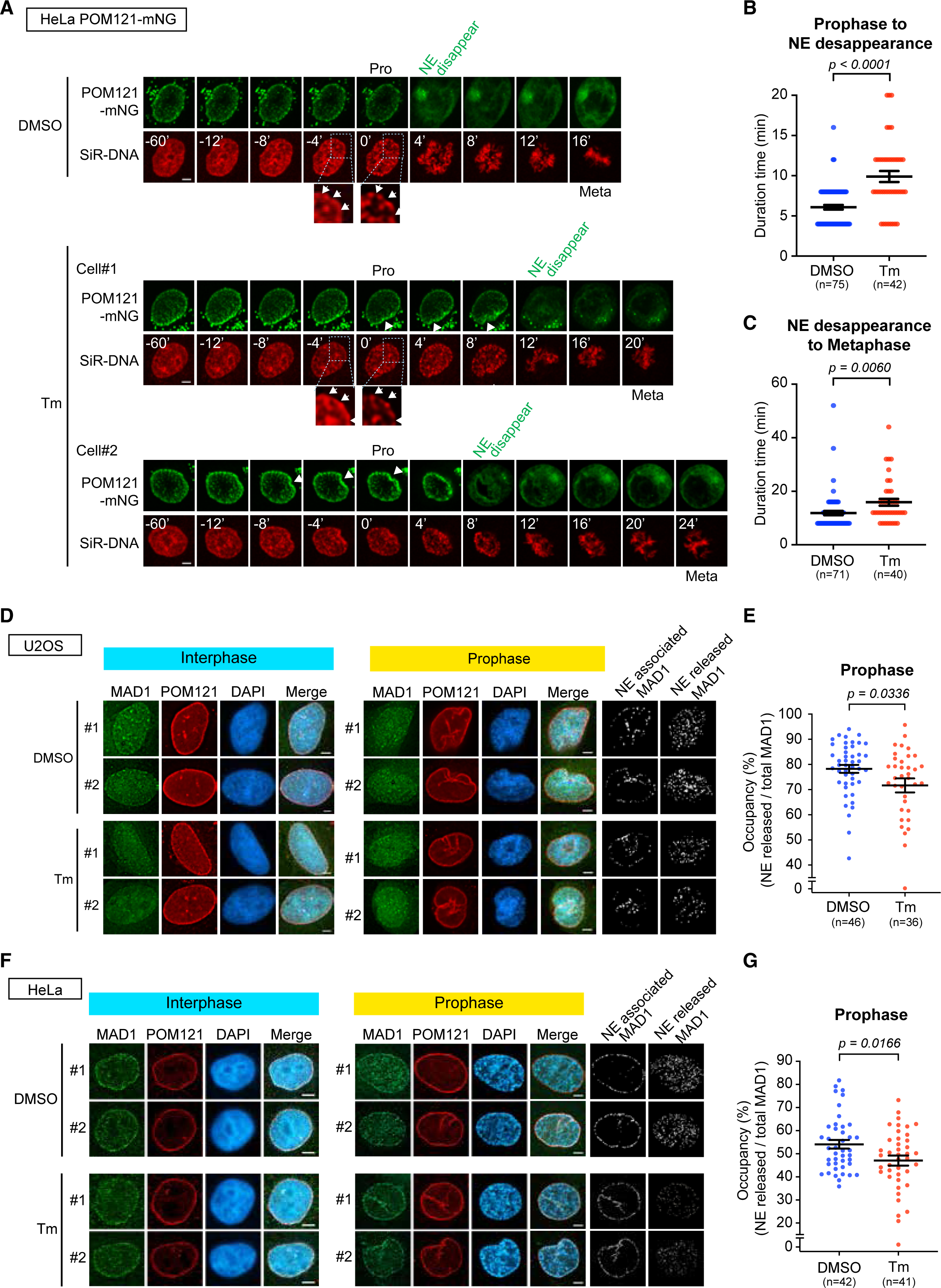
ER stress delays NEBD and impairs release of NE-localized MAD1 during mitotic entry. **(A)** Time-lapse imaging of mitotic entry in HeLa cells stably expressing POM121–mNeonGreen (mNG) under control or ER stress conditions. Cells were stained with SiR–DNA, pretreated with DMSO or Tm; (10 µg/mL) for 6 h, and imaged every 4 min for 10 h. Prophase onset was scored by the transition from continuous rim (interphase) to punctate/condensed SiR–DNA (magnified images of SiR-DNA for Prophase (0’) and 4 min prior to time 0’) and defined as time 0 min, and nuclear envelope (NE) breakdown was scored by loss of a continuous POM121–mNG rim signal; scale bars, 5 µm. (**B, C**) The timing of NE remodeling was quantified by measuring the duration from prophase onset to NE disappearance (B) and from NE disappearance to metaphase (C) in DMSO- and Tm-treated cells. These intervals report how ER stress alters the kinetics of NEBD and subsequent progression into metaphase. (**D–G**) MAD1 colocalizes with POM121 at the NE during interphase and prophase in both U2OS (**D, E**) and HeLa (**F, G**) cells and is released from the NE as cells commit to mitosis. Binarized MAD1 and POM121 images were used to distinguish NE-associated (POM121-colocalized) versus NE-released (non-NE) MAD1 and to measure the fraction of the NE region occupied by MAD1 signal in interphase and prophase. Scale bars, 5 µm. P values were calculated using Student’s t test, and data in (**E**) and (**G**) are presented as mean ± SEM from individual cells.

In DMSO-treated controls, the NE often exhibited increased wrinkling shortly before NE disappearance—defined as the time when POM121–mNG fluorescence began to diffuse beyond the nuclear region—consistent with prophase NE invagination (PNEI) described previously (Hebbar et al., 2008) and closely followed by NE breakdown during prometaphase (**Figure 5A**). In contrast, Tm–treated cells exposed to ER stress showed altered NE dynamics: although PNEI still occurred, the NE persisted for an extended period, and NE disappearance was substantially delayed (**Figure 5A–C**). Similar kinetic differences in NE disassembly were observed using Nup153 as an independent NPC marker (***Figure S4A***).

Microtubule attachment to sister chromatids requires the assembly of kinetochore proteins at centromeres, and several of these proteins localize to the NE prior to mitosis. During early mitosis, NEBD releases NE-associated kinetochore factors, allowing their translocation to kinetochores and promoting proper spindle attachment (Alfonso-Perez et al., 2019; Rasala et al., 2006; Zuccolo et al., 2007).. To determine whether ER stress perturbs this step, we focused on the spindle checkpoint protein Mitotic Arrest Deficiency 1 (MAD1), which is required for accurate chromosome alignment and for monitoring microtubule attachment (Akera et al., 2015; Lara-Gonzalez et al., 2021a; London and Biggins, 2014). Previous studies have shown that MAD1 dissociates from the NE into the nucleoplasm at early prophase, at a time when nuclear rim staining of nuclear basket components such as Drosophila Megator or human Tpr remains detectable (Cunha-Silva and Conde, 2020; Mossaid et al., 2020). To evaluate whether ER stress affects MAD1 release, we performed immunofluorescence using anti-MAD1 and anti-POM121 antibodies and analyzed cells at prophase, when chromosome condensation had initiated but the NE remained intact (**Figure 5D–G**).

We quantified MAD1 distribution between the NE and nucleoplasm. MAD1 fluorescence intensity was measured in regions of interest within the nucleoplasm, and NE-associated MAD1 was defined as MAD1 signal colocalizing with POM121 (see Materials and Methods for details). In ER-stressed U2OS cells, MAD1 signal remained enriched at POM121-positive NE regions, with correspondingly reduced MAD1 intensity in the nucleoplasm compared to DMSO controls (**Figure 5D–E**). This pattern indicates that ER stress delays MAD1 release from the NE, potentially compromising timely kinetochore loading and chromosome alignment. A similar delay in MAD1 redistribution was observed in HeLa cells (**Figure 5F–G**). These observations suggest that ER stress impedes MAD1 release by prolonging NE.

### Mitotic arrest by MG132 remodels ER architecture

Our findings establish that ER and nuclear envelope (NE) structural homeostasis are intricately linked to mitosis beyond their established roles in DNA- or genome-associated processes. The metaphase-to-anaphase transition represents a major cell cycle checkpoint, where perturbations—including non-degradation of Cyclin B, delayed spindle checkpoint silencing, Cdc20 depletion, or CENP-E inhibition—prolong mitotic progression (Nasmyth and Haering, 2009; Stevens et al., 2011). To determine whether such anaphase delays independently perturb ER homeostasis, apart from ER stress–induced chromosome misalignment, we treated cells with a low concentration (0.2 μM) of the proteasome inhibitor MG132 (Jia et al., 2011; Min and Lindon, 2012). This dose prolonged metaphase duration without inducing chromosome misalignment or spindle defects (**Figure 6A–B**, lanes 5 vs 6; ***Figure S5A***, lanes 1–2), consistent with prior reports (Uetake and Sluder, 2010). Higher MG132 concentrations, by contrast, cause pronounced metaphase arrest (Daum et al., 2011; Potapova et al., 2006; van Heesbeen et al., 2014).

**Figure 6.**
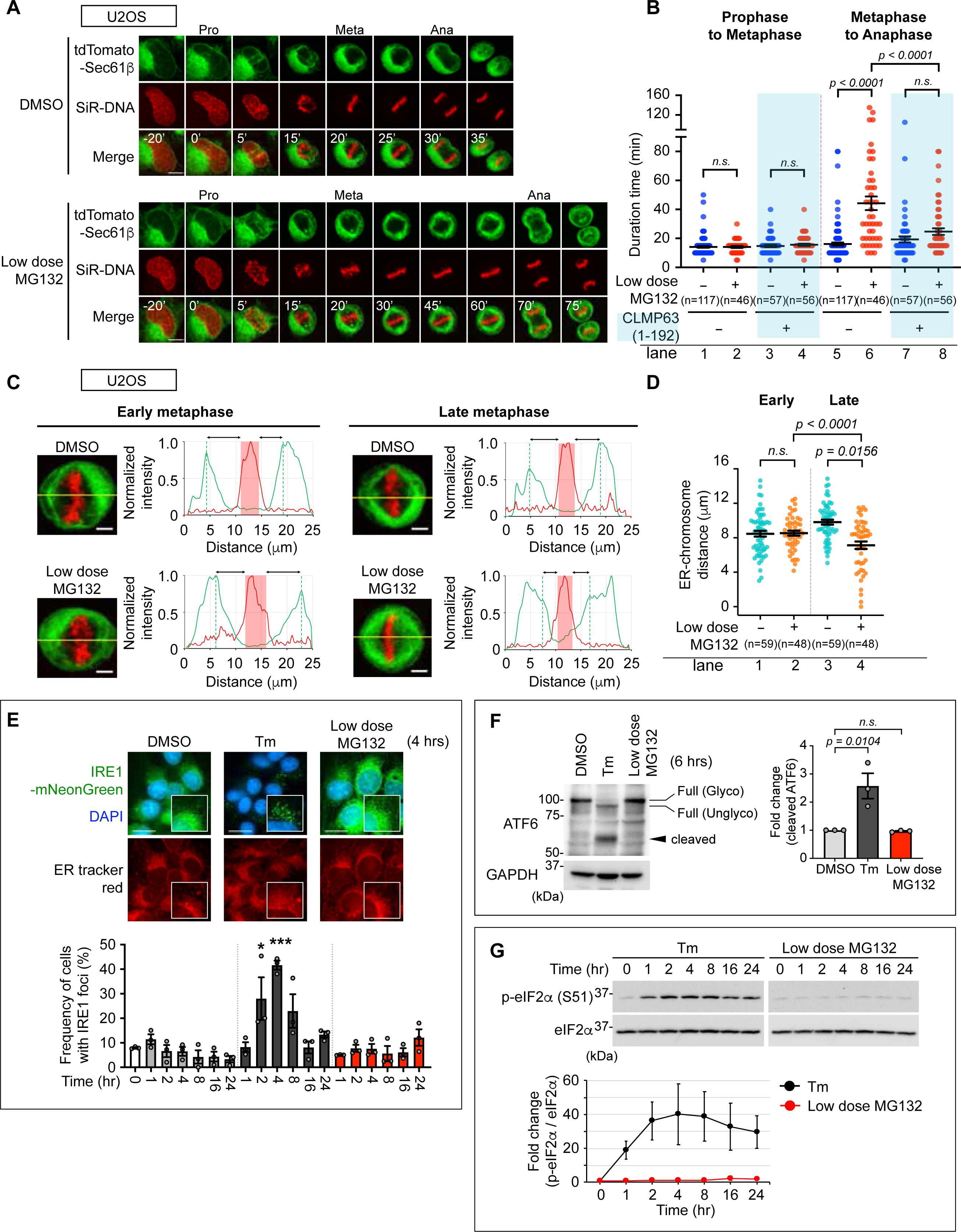
Extending metaphase by dampening APC/C-dependent proteolysis destabilizes ER organization and modestly engages the UPR. (**A**) Time-lapse imaging of U2OS cells stably expressing GFP–Sec61β reveals that low-dose MG132 (0.2 µM) prolongs metaphase while still permitting anaphase onset, creating a window to visualize how the ER remodels under sustained metaphase tension. Prophase onset, marked by chromosome condensation, was designated as time 0 min. Scale bars, 10 µm. (**B**) Quantitative analysis of mitotic phase durations under these conditions shows that even partial inhibition of proteasome-dependent APC/C activity is sufficient to extend metaphase. U2OS cells expressing GFP–Sec61β or CLIMP63(1–192) were pretreated with DMSO or MG132 (0.2 µM) as indicated. Data represent mean ± SEM from individual cells. (**C**) Representative images of early and late metaphase (as defined in Figure 1) illustrate that ER–chromosome relationships become progressively distorted when metaphase is prolonged. Corresponding line-scan profiles of normalized GFP–Sec61β (green) and SiR–DNA (red) intensities highlight a loss of the tight ER association with condensed chromosomes seen in control cells. Scale bars, 5 µm. (**D**) Quantification of ER-chromosome distances in cells expressing GFP–Sec61β demonstrates that low-dose MG132 consistently decreased ER-chromosome distances, linking altered ER architecture to a defined change in mitotic timing. Data represent mean ± SEM from individual cells. (**E**) To determine whether this mitotic perturbation engages ER stress signaling, IRE1 activity was monitored by IRE1–mNeonGreen oligomerization. Cells treated with DMSO, Tm, or MG132 for the indicated times show an increase in the fraction of cells with IRE1 foci under Tm but not by MG132. Scale bars, 20 µm. (**F**) ATF6 pathway engagement was assessed in parallel by immunoblotting for ATF6 cleavage. U2OS cells treated with DMSO or MG132 (0.2 µM) for 6 h did not exhibit a detectable increase in the cleaved (∼50 kDa) ATF6 fraction relative to full-length (∼100 kDa) species. Tm treatment generated cleaved ATF6, showing the robust activation as expected. Fold changes in cleaved ATF6 were normalized to DMSO controls, with GAPDH as a loading control. (**G**) PERK signaling branch activation under low-dose MG132 was evaluated by monitoring eIF2α phosphorylation at Ser51. MG132 (0.2 µM) did not induce a time-dependent increase in phospho-eIF2α while Tm (10 µg/mL) treatment induced robust levels of phospho-eIF2α. Phospho-eIF2α levels were normalized to total eIF2α and expressed relative to time 0, and P values were determined by one-way ANOVA with Sidak’s multiple comparisons test. In (B) and (D), data represent mean ± SEM from individual cells from three independent time lapse experiments; in (E–G), mean ± SEM from three independent experiments

We assessed whether this metaphase prolongation altered ER–chromosome organization using line-plot analyses (**Figure 6C–D**). In early metaphase, ER–chromosome spacing remained consistent between DMSO- and MG132-treated cells. Strikingly, however, prolonged late metaphase under MG132 caused a significant reduction in ER–chromosome distance (**Figure 6C–D**, lanes 3–4), indicating that extended metaphase duration alone—independent of ER stress—disrupts ER architecture relative to metaphase DNA.

Since proteasome inhibitors, at certain doses, might trigger ER stress by causing the accumulation of unfolded proteins in the ER, we assessed UPR activation at this lower concentration of MG132 contributed for the observed metaphase extension and ER-metaphase chromatin spacing. IRE1 oligomerization was measured by IRE1-mNeonGreen foci formation (**Figure 6E**), and activation of the other UPR branches, PERK and ATF6α, was evaluated in U2OS cells (**Figure 6F-G**) (Belyy et al., 2020; Haze et al., 1999). Thus, our results are in agreement with earlier reports where 0.2 μM MG132 did not activate IRE1, PERK, or ATF6α, unlike ER stress induced from Tm (Kroiss et al., 2016; Wang et al., 2019). Taken together, these results revealed that MG132 at this dosage prolonged metaphase without inducing ER stress or chromosome assembly defects (**Figure 6A–B, lanes 5 vs 6**).

Finally, to further dissect the role of ER structure, we tested overexpression of the ER-shaping mutant CLIMP63(1–192). As observed under ER stress, CLIMP63(1–192) counteracted MG132-induced ER deformation, restored metaphase delay (**Figure 6B**, lane 6 vs 8, and ***Figure S5B***) and ER–chromosome spacing (***Figure S5C-D,*** lanes 3–4 vs 7–8 and ***Figure S5E–F***). These results demonstrate that ER remodeling is exquisitely sensitive to mitotic timing perturbations driven by MG132-mediated APC/C inhibition and accumulation of substrates such as securin and cyclins.

Together, these data reveal a bidirectional interplay: prolonged metaphase remodels ER organization, which in turn exacerbates mitotic delay, suggesting a positive feedback loop. ER homeostasis thus emerges as a dynamic integrator of organelle structure and chromosome segregation fidelity during stressed mitosis.

## Discussion

In this study, we identify a previously underappreciated functional link between the ER and the NE—two structurally continuous yet functionally distinct membrane systems—as a critical determinant of the mammalian cell cycle. By examining mammalian cells undergoing open mitosis under ER stress, we find that disruption of ER homeostasis impairs mitotic progression and causes a specific delay in the metaphase–anaphase transition. This delay coincides with marked ER disorganization and defects in early mitotic events, including centrosome separation, spindle assembly, and chromosome alignment (**Figure 7**). The metaphase–anaphase delay induced by ER stress is attributable, in part, to delayed NE breakdown, which postpones the release of MAD1, an NE-associated kinetochore protein, due to disruption of the ER–NE interface. Overexpression of ER-shaping proteins such as REEP4 and a truncated form of CLIMP63 (1–192) mitigates these defects, indicating that proper ER–NE coupling is required to maintain mitotic fidelity beyond the level of spindle assembly.

**Figure 7.**
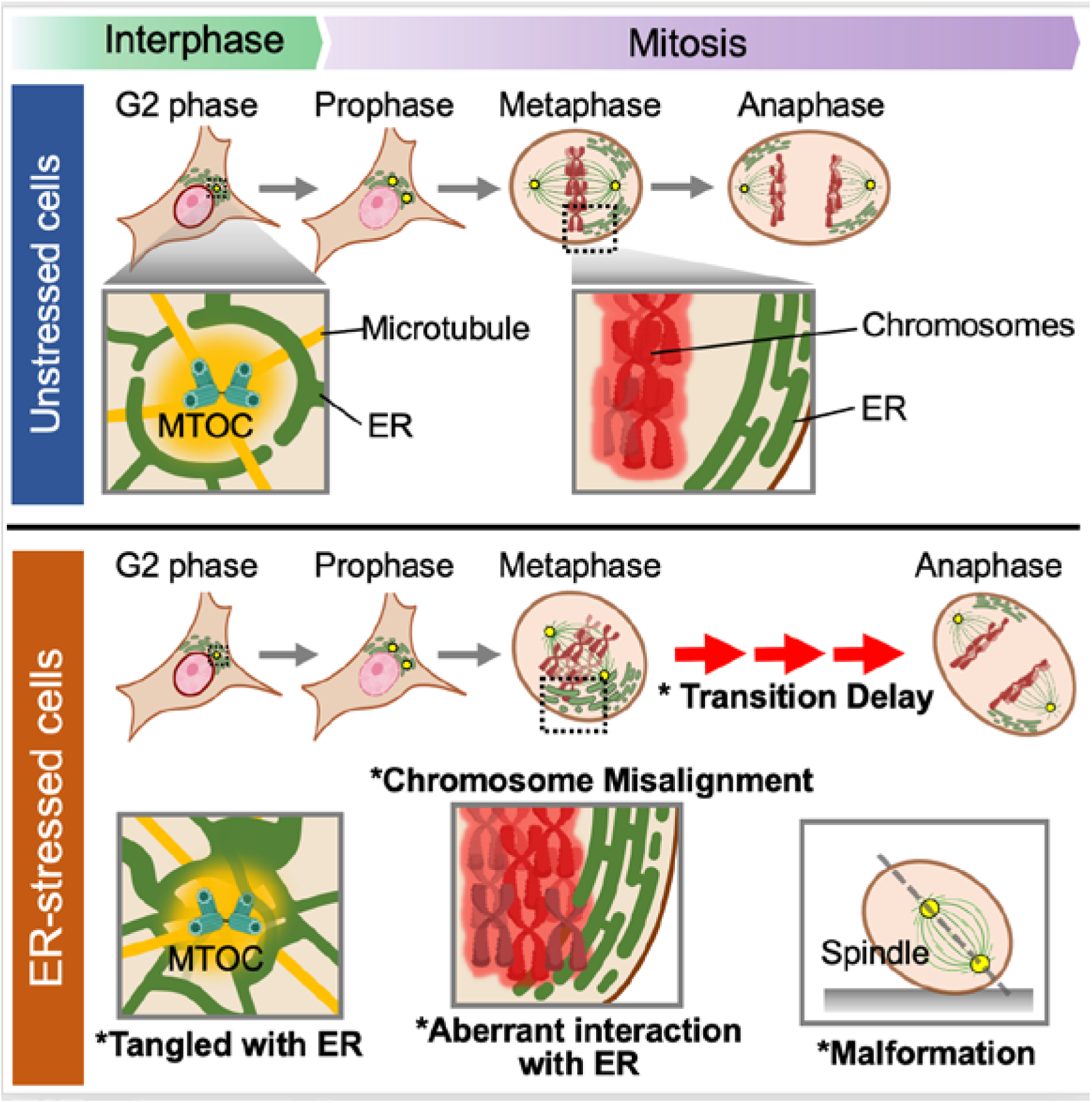
Integrated model linking ER organization to mitotic structure and progression. Disruption of ER homeostasis or induction of ER stress perturbs ER organization at microtubule-organizing centers (MTOCs) during G2, compromising the centriculum and associated ER networks. During metaphase, ER-stressed cells exhibit abnormal ER–chromosome spatial relationships, leading to chromosome misalignment and spindle defects that collectively prolong the metaphase-to-anaphase transition. In parallel, experimentally prolonging metaphase with low-dose MG132 alters ER–chromosome organization even in the absence of overt ER stress, indicating that mitotic timing itself feeds back on ER architecture. Together, these findings support a model in which precise spatial positioning of the ER relative to chromosomes is not merely permissive but functionally integrated into faithful mitotic progression.

Disruption of ER homeostasis also perturbs tubulin organization early in the cell cycle. Although centrosomes are traditionally regarded as membrane-less organelles, they are closely enveloped by ER membranes that may impart lipid bilayer–like properties and influence their structural stability. Under ER stress, these pericentrosomal membranes lose their ordered organization, contributing to spindle defects and the observed metaphase delay. Restoration of ER structural homeostasis by CLIMP63(1–192) expression corrected these abnormalities, underscoring that an intact ER lipid network is required early in mitosis to position centrosomes correctly at the outer nuclear membrane as components of an integrated membrane system rather than as autonomous entities.

Entry into anaphase marks one of the most decisive transitions in the cell cycle. When its molecular choreography falters, chromosome segregation is compromised, giving rise to aneuploidy—a defining feature of tumorigenesis (Lara-Gonzalez et al., 2021b). Seeking to probe how mitotic timing intersects with organelle architecture, we asked whether anaphase entry delays, even those uncoupled from spindle defects, might reshape the organization of the ER and NE. In line with this idea, delaying mitotic progression with the proteasome inhibitor MG132—which blocks APC/C-mediated degradation of regulators such as cyclin B and securin (Yamano, 2019)—disrupted ER–NE architecture and altered their coordination during mitosis. Intriguingly, overexpression of CLIMP63(1–192) alleviated this extended metaphase–anaphase delay, implying that structural features of the ER can influence mitotic timing itself. Although high doses of MG132 are known to impair ER proteostasis and arrest cells in metaphase, the low-dose treatment used here permitted entry into anaphase without activating the unfolded protein response. Thus, the observed delay was not a direct consequence of ER proteostatic stress. Instead, a modest prolongation of mitosis—independent of immediate ER stress—was sufficient to remodel ER organization and positioning. These findings reveal that the architecture of the ER–NE system is intimately tuned to mitotic timing through a mechanism that operates largely outside canonical ER proteostasis pathways.

Mitotic remodeling of the ER involves its redistribution toward the cell periphery and the formation of a central ER exclusion zone (Carlton et al., 2020; Lu et al., 2009). Our live-cell imaging shows that ER stress disrupts this process, causing aberrant ER accumulation around chromosomes and delaying anaphase onset. Cytoskeletal forces and their coupling to the ER via the LINC complex and ER-shaping proteins govern organelle positioning during mitosis (Jongsma et al., 2015; van Bergeijk et al., 2016). Our results suggest that ER homeostasis is critical for proper LINC complex positioning and that ER stress–induced defects, or alterations in LINC components, may disrupt this coordination, leading to changes in spindle size or orientation and ultimately prolonging the metaphase–anaphase transition. Indeed, recent work has also linked perturbations of the LINC complex to UPR activation (Belaadi et al., 2022; Buchwalter et al., 2019; Turgay et al., 2014), supporting a bidirectional relationship between NE–cytoskeletal coupling and ER stress signaling. Another, non-exclusive, possibility is that ER stress increases overall ER mass through activation of UPR pathways (Moncan et al., 2021). Excessive ER expansion has been shown to promote abnormal ER–chromosome associations that impair chromosome alignment during mitosis (Merta et al., 2021), and aberrant or enforced ER–chromosome contacts are increasingly recognized as a barrier to proper chromosome alignment (Champion et al., 2019; Ferrandiz et al., 2022). Together, these observations support dual mechanisms—disrupted cytoskeletal coupling and ER overexpansion—by which ER stress impairs mitotic ER organization and delays faithful chromosome segregation.

Additionally, ER morphology undergoes dynamic remodeling during mitosis. Under normal conditions, the ER transitions to a tubule-dominant network, whereas ER stress biases it toward sheet formation through enhanced lipid synthesis and ER expansion (Chen et al., 2023; Corazzari et al., 2017; Kwon et al., 2023). REEP4, a tubule-promoting protein, supports the high membrane curvature characteristic of this mitotic transition and has been implicated in establishing metaphase “ER clearance” away from chromatin (Kumar et al., 2019; Schlaitz et al., 2013). Consistent with these functions, REEP4 overexpression alleviated ER stress–induced metaphase delay in our system, reinforcing the importance of ER organization for mitotic timing. However, the relationship between ER morphology and mitotic progression appears more complex than a simple sheet-versus-tubule dichotomy. Overexpression of full-length CLIMP63, a sheet-promoting protein, did not rescue metaphase delay to the same extent as REEP4 (data not shown), whereas the truncated variant CLIMP63(1–192) restored both mitotic timing and chromosome alignment. This is consistent with ultrastructural analyses showing that CLIMP63(1–192) promotes narrow-lumen, high-curvature ER morphologies typical of tubular networks (Shen et al., 2019; Wang et al., 2022; Xu et al., 2023). Alternatively, because CLIMP63(1–192) retains the cytosolic and transmembrane domains and can interact with endogenous CLIMP63, it may function as a dominant-negative, attenuating the sheet-promoting activity of full-length CLIMP63 and thereby favoring a tubule-dominant ER organization during mitosis. Thus, CLIMP63(1–192) may shorten metaphase either by enhancing high-curvature membrane geometry or by antagonizing endogenous CLIMP63, shifting the ER toward a tubule-rich state that supports efficient chromosome alignment and anaphase onset. Together, these findings position ER-shaping proteins as active determinants of mitotic efficiency under stress conditions.

Our results revealed that ER stress accelerates separation of α-tubulin–marked MTOC/centrosomal foci while simultaneously delaying NEBD and bipolar spindle assembly, consistent with premature centrosome separation uncoupled from proper maturation. Proper centrosome function depends on maturation of the pericentriolar material (PCM) and full MTOC activity before centrosome separation. These findings therefore suggest that, under ER stress, centrosomes may separate before achieving full maturity. Duplicated centrosomes are normally tethered by a linker that is degraded only after centrosome maturation, through the coordinated action of mitotic kinases (PLKs and Aurora kinases) and Eg5-dependent motor forces (Raaijmakers et al., 2012). Given that the unfolded protein response (UPR) broadly perturbs upstream signaling pathways that regulate centrosome maturation, including AKT and MAPK (Chen et al., 2023; Hetz et al., 2020), ER stress could disturb these regulatory inputs and thereby promote precocious centrosome separation, compromising orderly mitotic progression and increasing the likelihood of downstream mitotic errors.

Furthermore, under unstressed conditions, the ER adopts a pericentrosomal, fenestrated organization before mitotic entry, with membranes surrounding but not directly overlapping the centrosomal region, such that the MTOC appears to reside within an “opening” of the ER network. Under ER stress, this architecture is lost: centrosomal foci become entangled with ER membranes, and the spatial segregation between the MTOC and surrounding ER is no longer apparent. Notably, in Drosophila, ER disruption inhibits centrosome maturation and induces spindle malformation concomitant with abnormal ER dynamics (Araujo et al., 2023; Rollins and Blankenship, 2023). One mechanistic explanation involves the “centriculum,” an ER-derived structure described in *C. elegans* and Drosophila that forms in close association with the centrosome and serves as a transient platform regulating PCM size and microtubule nucleation (Maheshwari et al., 2023; Rollins and Blankenship, 2023). By analogy, disruption of ER organization in mammalian cells could compromise a centriculum-like arrangement, thereby impairing centrosome maturation and spindle assembly. Future work will be required to test this possibility directly.

Chromosome misalignment, manifested as broadened metaphase plates under ER stress, further underscores the contribution of ER morphology to mitotic fidelity. Proper plate formation requires robust mechanical tension between sister chromatids, generated by spindle microtubules and centromeric cohesion complexes (Jaqaman et al., 2010). Perturbations in cohesion, condensin, or CTCF function are known to broaden metaphase plates by weakening inter-chromatid tension (Ribeiro et al., 2009; Toyoda and Yanagida, 2006; Walsh and Stephens, 2025), and ER stress likely adds a spatial, organelle-based dimension to this defect. In our system, ER stress was accompanied by increased ER–chromosome overlap, particularly at the edges of the metaphase plate, suggesting that mispositioned ER physically or sterically interferes with chromosome compaction and the establishment of proper tension. Such aberrant ER–chromosome contacts could hinder stable kinetochore–microtubule attachments and efficient error correction, thereby promoting defective alignment and plate broadening.

Our previous work in yeast identified the ER stress surveillance (ERSU) pathway, which halts cell cycle progression when ER integrity is compromised (Babour et al., 2010; Chao et al., 2019; Pina et al., 2018; Pina et al., 2016; Pina and Niwa, 2015). Several ERSU components—including SLT2, septins, and the early sphingolipid intermediate phytosphingosine (PHS)—are conserved in mammals, suggesting the existence of a related surveillance mechanism, even if the molecular details may differ. Oscillatory ERK signaling contributes to spindle assembly (Horne and Guadagno, 2003; Roberts et al., 2002), and ER stress perturbs this dynamic (Galan et al., 2014; Hwang et al., 2013; Kato et al., 2014). In parallel, ER stress elevates sphingolipid intermediates such as dihydrosphingosine (DHS) and dihydroceramide (DHC), mammalian equivalents of PHS, which bind the ATF6 transmembrane domain, activate ATF6, and drive ER biogenesis (Tam et al., 2018). These observations support a model in which ERK-, septin-, and sphingolipid-dependent signaling modules form components of a mammalian ER stress–responsive mitotic surveillance system that integrates ER functional homeostasis with accurate and timely chromosome segregation. Future work will uncover a mechanism corresponding like ERSU pathway in yeast regulates ER stress-induced mitotic events that we described here.

Metaphase delay serves as a major checkpoint-like pause to prevent diverse mitotic errors, including—but not limited to—those arising from ER stress–induced spindle assembly defects. In this context, altered APC/C-dependent proteolysis by low-dose MG132 was accompanied by deformed ER organization around the metaphase chromosomes and MTOC/centrosomes, further highlighting the intimate relationship between ER architecture and mitotic timing. Together, these findings indicate that ER homeostasis acts as an essential integrator of accurate and timely mitotic events and reinforce the emerging view that the ER functions not as a passive bystander during cell division, but as an active structural and signaling hub whose integrity is crucial for faithful mitotic progression. Future studies will be required to define the molecular machinery that regulates mitotic ER remodeling and to dissect how it interfaces with canonical cell cycle checkpoints.

## Supplemental figure legends

***Figure S1*. ER stress delays anaphase onset and disrupts ER organization**

(**A**) Time-lapse imaging of mitotic progression under ER stress. HeLa cells stably expressing GFP–Sec61β and H2B–mCherry were pretreated with DMSO, tunicamycin (Tm; 5 µg/mL, 6 h), or β-mercaptoethanol (β-ME; 2.5 mM, 2 h), and imaged every 5 min for 10 h, with prophase onset defined as time 0 min. Scale bar, 10 µm.

(**B**) Quantification of mitotic phase durations under the conditions in (A) shows that ER stress prolongs the metaphase-to-anaphase transition. Data represent mean ± SEM from individual cells.

(**C**) Representative images and line-scan analyses of ER–chromosome organization in early and late metaphase HeLa cells. Early and late metaphase are defined in Figure 1. Normalized fluorescence intensities of ER (GFP–Sec61β, green) and chromosomes (H2B–mCherry, red) were plotted along the yellow line shown in the images. Chromatin regions (light red shading) and the nearest ER fluorescence peaks (green dashed lines) were used to measure ER–chromosome distances.

(**D**) Quantification of ER–chromosome distances in DMSO-, Tm-, or β-ME–treated HeLa cells at early and late metaphase, revealing increased ER–chromosome separation under ER stress. Data represent mean ± SEM from individual cells.

(**E**) Analysis of ER overlap at the ends of “metaphase” chromosomes in U2OS cells. Binary images generated from ER and DNA signals were used to define the upper and lower 20% of the metaphase chromosome length as chromosome ends, and ER–DNA overlapping and DNA end areas were measured to calculate ER occupancy.

(**F, H**) Representative enlarged images showing ER overlap at the ends of “metaphase” chromosomes in DMSO-, Tm-, or Tg–treated U2OS cells (F) and DMSO-, Tm-, or β-ME–treated HeLa cells (H). DNA regions are outlined in red, and overlapping ER regions are shown in white.

(**G, I**) Quantification of ER overlap at metaphase chromosome ends in DMSO-, Tm-, or Tg–treated U2OS cells (G) and DMSO-, Tm-, or β-ME–treated HeLa cells (I), demonstrating reduced ER occupancy at Chromosome ends under ER stress exhibit reduced ER occupancy, consistent with compromised ER–chromosome coupling at metaphase. Statistical analysis for (B), (D), (G), and (I) was performed using one-way ANOVA with Sidak’s multiple comparisons test, and data represent mean ± SEM from individual cells.

***Figure S2*. Overexpression of CLIMP63(1–192) and REEP4 mitigates ER stress–induced mitotic defects**

(**A**) ER association with chromosome termini in early and late “metaphase” U2OS cells overexpressing CLIMP63(1–192) is visualized in enlarged views, which highlight chromosome ends with DNA boundaries outlined in red and overlapping ER signal shown in white.

(**B**) Quantification of ER overlap at early versus late “metaphase” chromosome ends in DMSO- and tunicamycin (Tm)-treated U2OS cells, as shown in (A), reveals that CLIMP63(1–192) overexpression partially restores ER coverage at chromosome termini under ER stress.

(**C**) Representative images and line-scan analyses of ER–chromosome distributions in early and late metaphase HeLa cells overexpressing REEP4 under Tm treatment show that enhanced REEP4 levels help maintain a closer ER–chromosome apposition. Normalized fluorescence intensities of ER (GFP–REEP4, green) and chromosomes (NucBlue, red) are plotted along the indicated line. Scale bars, 5 µm.

(**D**) Quantification of ER–chromosome distances under the conditions shown in (C), compared with cells lacking REEP4 overexpression, demonstrates that REEP4 overexpression reduces ER–chromosome separation in Tm-treated cells. P values in (B) and (D) were obtained by one-way ANOVA followed by Sidak’s multiple comparisons test, and data represent mean ± SEM from individual cells.

**Figure S3. ER stress disrupts spindle architecture and centrosome integrity.**

(**A**) Representative time-lapse images show MTOC separation and spindle formation in cells subjected to ER stress, progressing to prophase (time 0 min, defined by the appearance of condensed chromatids) through subsequent mitotic stages. Scale bars, 10 µm.

(**B–D**) Quantitative analysis of metaphase spindle architecture in HeLa cells under ER stress. Spindle length and width (B), spindle angle (C), and cell size (D) were measured in HeLa cells stably expressing GFP–α-tubulin, stained with SiR–DNA, and pretreated with DMSO, Tm; (5 µg/mL, 6 h), or β-ME; (2.5 mM, 2 h) before imaging. Time-lapse microscopy was performed at 5-min intervals, and metaphase spindle images were collected one frame before anaphase onset. P values were calculated using one-way ANOVA followed by Sidak’s multiple comparisons test. Mean values were derived from three independent experiments, with data shown as mean ± SEM from individual cells. Scale bars, 5 µm.

**Figure S4. ER stress delays nuclear envelope breakdown (NEBD) and MAD1 release.**

(**A**) Representative time-lapse images show mitotic progression in U2OS cells stably expressing GFP–Nup153 under conditions of ER stress. Cells were labeled with SiR–DNA and pretreated with either DMSO or Tm (10 µg/mL, 6 hours), and imaged every 4 minutes for 10 hours, with the onset of prophase designated as time 0 minutes. Scale bars, 10 µm.

***Figure S5*. Overexpression of ER protein CLIMP63(1–192) mitigates MG132-induced metaphase extension and re-stores ER–chromosome organization.**

(**A**) Metaphase chromosome alignment in U2OS cells treated with low-dose MG132 (0.2 µM). Metaphase chromosomes were classified as in Figure 1C: fully aligned (blue), broad metaphase plate (green), no or broad metaphase plate (purple), and scattered plate (red). Spindle width (maximum microtubule bundle span) was quantified from GFP–α-tubulin time-lapse images acquired one frame before anaphase.

(**B**) Representative time-lapse images of U2OS cells stably expressing mCherry–CLIMP63(1–192) under low-dose MG132 (0.2 µM). Cells were labeled with SiR–DNA and pretreated with DMSO or MG132 for 2 hours, followed by imaging every 5 minutes for 10 hours. Scale bars, 10 µm.

(**C**) ER–chromosome spatial relationships at early and late metaphase in CLIMP63(1–192)–expressing cells treated with MG132. Normalized fluorescence intensities of ER (mCherry–CLIMP63(1–192), green) and chromosomes (SiR–DNA, red) are plotted, revealing that CLIMP63(1–192) expression helps maintain ER proximity to metaphase chromosomes under proteasome inhibition. Scale bars, 5 µm.

(**D**) Quantification of ER–chromosome distances in U2OS cells treated with or without MG132, comparing CLIMP63(1–192)–nonexpressing cells (lanes 1–4, from Figure S5D) and CLIMP63(1–192)–expressing cells (lanes 5–8). CLIMP63(1–192) overexpression reduces MG132-induced increases in ER–chromosome separation.

(**E**) Overlapping ER signal at metaphase chromosome ends in early and late metaphase U2OS cells overexpressing CLIMP63(1–192). Enlarged insets show chromosome ends with DNA regions outlined in red and overlapping ER regions highlighted in white in binary images.

(**F**) Quantification of overlapping ER regions at metaphase chromosome ends in cells treated with DMSO or MG132 indicates that CLIMP63(1–192) overexpression partially restores ER occupancy at chromosome termini under MG132 treatment. P values for (A), (D), and (F) were calculated using one-way ANOVA with Sidak’s multiple comparisons test, and data represent mean ± SEM from individual cells

## Materials and Methods

### Cell Culture

U2OS cells were maintained in McCoy’s 5A medium (Cat# 16600082; Gibco, Waltham, MA, USA) supplemented with 10% fetal bovine serum (FBS; Cat# A52567-01; Gibco). HeLa cells and Lenti-X 293T cells were maintained in DMEM (Cat# 10-013-CV, Corning, Corning, NY, USA) supplemented with 10% FBS. U2OS cells were a gift from Prof. Martin Hetzer (Institute of Science and Technology Austria, Austria). HeLa cells stably expressing GFP-Sec61β/H2B-mCherry and HeLa cells stably expressing GFP-Tubulin were gifts from Prof. Katharine Ullman (The University of Utah, USA). Lenti-X 293 cells were a gift from Prof. Matthew Dougherty (University of California San Diego, USA). U2OS cells expressing IRE1-mNeonGreen (mNG) under the control of a tetracycline-inducible promoter (Belyy et al., 2020) were a gift from Prof. Peter Walter (University of California San Francisco, USA). All cells were maintained at 37 °C in a humidified incubator with 5% CO2.

Cells were routinely checked for mycoplasma contamination using PCR or DNA staining methods. For the DNA staining method, fixed cells were stained with 5 μg/mL 4, 6-diamidino-2-phenylindole-dihydrochloride (DAPI; Cat# 46190; Thermo Fisher Scientific, Waltham, MA) and verified to have no cytoplasmic fluorescence under a fluorescence microscope. For the PCR methods, cell culture supernatant was collected from semi-confluent cells cultured for three days without medium change. After removal of debris by centrifugation, the supernatant was directly used for nested PCR with the following primers: Fwd_1st: 5’-ACACCATGGGAGCTGGTAAT-3’, Rv_1st: 5’-CTTCATCGACTTTCAGACCCAAGGCAT-3’, Fwd_nested: 5’-GTTCTTTGAAAACTGAAT-3’, Rv_nested: 5’-GCATCCACCAAAAACTCT-3’

### Plasmids

L304-EGFP-Tubulin-WT (Addgene plasmid #64060), mCherry-CLIMP63(1-192) (Addgene plasmid #186955), pAcGFP-C1-Sec61β (Addgene plasmid #62008), and pEGFP(C3)-Nup153 (Addgene plasmid #64268) were obtained from Addgene (Watertown, MA, USA). pLEX307-APN-puro (Addgene plasmid #158454), pLenti-CMV-GFP Hygro (656-4) (Addgene plasmid #17446), psPAX2 (Addgene plasmid #12260), and pMD2.G (Addgene plasmid #12259) were gifts from Prof. Dougherty. mEmerald-Sec61β was a gift from Prof. Craig Blackstone (Massachusetts General Hospital). REEP4 cDNA was synthesized and cloned into pcDNA3.1+N-eGFP by GenScript Biotech (Piscataway, NJ). ptdTomato-Sec61β was constructed by subcloning Sec61β into the ptdTomato-C1 vector (Cat# 632533; Clontech). Lentiviral vectors (pLEX-puro-GFP-Sec61β, pLEX-puro-tdTomato-Sec61β, pLEX-puro-POM121-mNG, pLenti-puro-EGFP-Tubulin) were constructed by subcloning genes of interest, fused to fluorescent proteins, into pLEX307-puro or pLenti-hygro backbones. All constructs were sequence-verified by whole plasmid sequencing (Plasmidsaurus, Eugene, OR, USA).

### Generation of Stable Cell Lines

U2OS cells stably expressing mEmerald-Sec61β, GFP-CLIMP63, mCherry-CLIMP63(1-192), GFP-REEP4, or GFP-RTN4 were generated by stable transfection. Cells were transfected with plasmids using Effectene Transfection Reagent (Cat# 301425; Qiagen, Hilden, Germany) according to the manufacturer’s protocol. Two days after transfection, cells were selected with 500 μg/mL G418 (Cat# 10131035; Gibco) for at least 5 days. Lentiviral supernatants were prepared by co-transfecting pLEX307-puro or pLenti-hygro vectors with psPAX2 and pMD2.G into Lenti-X 293T packaging cells at a ratio of 4:2:1. Twenty-four hours post-transfection, the culture medium was replaced, and cells were incubated for an additional 24 hours. Viral supernatants were then filtered through 0.45 µm PVDF syringe filters (Cat# 25-241; Genesee Scientific, El Cajon, CA, USA) and stored at 4 °C or −80 °C until use. For lentiviral transduction, viral supernatants were diluted 1:3 to 1:10 in culture medium containing 5 μg/mL polybrene (Cat# TR1003; Millipore Sigma, Burlington, MA, USA). Cells seeded on a 6-well plate were cultured in 1 mL of viral dilution for 6 hours, after which 1 mL of culture medium was added, and cells were incubated overnight. Cells were then selected with 1 μg/mL puromycin (Cat# A1113803; Gibco) or 200 μg/mL hygromycin (Cat# 10687010; Thermo Fisher Scientific) for at least 5 days.

### Live-cell Imaging

Cells were seeded in μ-Plate 24-well plates (Cat# 82426; ibidi, Gräfelfing, Germany) or μ-Plate 96-well plates (Cat# 89626; ibidi) and imaged with a 40× objective lens at 37 °C and 5% CO2 using a CQ1 spinning disk confocal system (Yokogawa Electric Corp., Tokyo, Japan). Chromosomes were visualized using either NucBlue Live ReadyProbes Reagent (Cat# R37605; Thermo Fisher Scientific) or SiR-DNA (Cat# CY-SC007; Spirochrome, Stein am Rhein, Switzerland). For U2OS cells with SiR-DNA staining, verapamil (Spirochrome) was co-incubated to inhibit dye efflux pathways and enable homogenous DNA staining (Lukinavicius et al., 2015). All experiments were performed under asynchronous conditions.

For experiments in Figure 1 (tunicamycin, Tm) and Figure 2, cells were seeded in 24-well assay plates at a density of 6.0 × 10^5^ cells in 1 mL per well and pre-cultured overnight. The medium was then replaced with fresh medium containing NucBlue dye (1 drop per 6 mL) and 10 μg/mL Tm (Cat# 654380; Sigma-Aldrich, St. Louis, MO, USA), and time-lapse imaging was initiated. Images were acquired at 1.4 µm z-steps for a total of 6 slices. Three to six fields per well were imaged for 16 hours at 5-minute intervals.

For other experiments using Tm or thapsigargin (Tg), cells were seeded in 96-well assay plates at a density of 15,000 cells in 300 μL per well and pre-cultured overnight. Cells were then incubated with 10 μg/mL Tm or 200 nM Tg (Cat# 586005; Calbiochem, La Jolla, CA, USA) in the presence of 50 nM SiR-DNA, and 1 μM verapamil (for U2OS cells). After 6 hours of incubation, time-lapse imaging was initiated. Images were acquired at 1.2 µm z-steps for a total of 7 slices. Two to five fields per well were imaged for 8-12 hours at 4-minute intervals (for Figure 5) or 5-minute intervals (for all other experiments). For experiments with β-mercaptoethanol (β-ME) or low-dose MG132, cells were seeded in 96-well assay plates at a density of 15,000 cells in 300 μL per well and pre-cultured overnight. Cells were then incubated with 2.5 mM β-ME (Cat# M3148; Sigma-Aldrich) or 0.2 μM MG132 (Cat# 508338; Thermo Fisher Scientific) in the presence of SiR-DNA dye, and 1 μM verapamil (for U2OS cells). After 2 hours of incubation, time-lapse imaging was initiated. Images were acquired at 1.2 µm z-steps for a total of 7 slices. Three to five fields per well were imaged for 8-12 hours at 5-minute intervals.

### Imaging Analysis of Live Cells

Microscope images were processed and analyzed using CellPathfinder (Yokogawa Electric Corp.) and Fiji Software (Schindelin et al., 2012). Maximum-intensity projection (MIP) images from live-cell imaging were generated using the three central z-slices of each cell for analysis. For analysis of mitotic duration, mitotic phases are defined by DNA morphology as follows: prophase, when DNA condensation first became detectable; metaphase, when chromosomes aligned at the equatorial plate; anaphase, when metaphase chromosomes began to separate; and telophase, when DNA decondensation was initiated.

Analysis of ER-chromosome distance and ER-chromosome overlap at metaphase chromosome ends was performed using MIP images taken from metaphase onset (early metaphase) and one frame before anaphase onset (late metaphase). Images were preprocessed with a Gaussian blur filter (σ = 1). For ER-chromosome distance analysis, fluorescent intensity profiles were obtained along a line drawn perpendicular to the metaphase chromosomes and normalized to their maximum and minimum intensities. The ER-chromosome distance was defined as the distance between chromosomes exhibiting normalized intensities greater than 50% of the maximum and the nearest ER signal peaks on both sides. Because ER peaks differed in height between the two sides, each peak was defined relative to its own maximum with a 50% threshold. For ER-chromosome overlap analysis, binary images of the ER and chromosomes were generated using the MaxEntropy or Otsu auto thresholding algorithms. Chromosome ends were defined as the upper and lower 20% of the total chromosome length based on the chromosome binary mask. The overlapping ER signal within these regions was calculated as the percentage of occupancy.

For spindle measurements, spindle length (pole-to-pole distance) and spindle width (the maximum width of microtubule bundles) were measured from MIP images acquired from late metaphase frames. For measurements of spindle angle with respect to the substrate, z-slices containing the center of spindle poles were defined by their maximum tubulin intensity. The angle was calculated using the following formula: θ = ArcSin(Z/A), where A is spindle length and Z is the z-axis distance between the two spindle poles.

Graphs represent mean values calculated from individual cell measurements pooled from at least three independent experiments.

### Immunofluorescent Staining and Imaging

Coverslips (Cat# 1943-10012A; 12 mm diameter, #1 thickness, Neuvitro Corp., Vancouver, WA, USA) were coated with 0.1 mg/mL poly-L-Lysine (Cat# P9155; Sigma-Aldrich, St.Louis, MO) in 24-well plates for 15 minutes, rinsed with PBS, and air-dried. Cells were then seeded onto the coated coverslips. After drug treatment, cells were fixed with 4% paraformaldehyde (PFA) for 15 minutes and permeabilized with 0.1% Tx-100 for 5 minutes at room temperature (RT). Following PBS washes, cells were incubated with blocking solution (5% BSA in PBS) for 1 hour. Primary antibody incubation was performed for 2 hours at RT or overnight at 4°C using the following primary antibodies: anti-MAD1 (1:100; Cat# sc-47746; Santa Cruz Biotechnology, Dallas, Tx, USA), anti-POM121 (1:200; Cat# 15645-1-AP; ProteinTech, Rosemont, IL, USA). After PBS washes, cells were incubated with secondary antibodies for 30 minutes at RT: goat anti-rabbit Alexa Fluor 568 (1:2000; Cat# A-11036; Thermo Fisher Scientific), goat anti-mouse Alexa Fluor 568 (1:2000; Cat# A-11031; Thermo Fisher Scientific). Coverslips were mounted on glass slides using Aqua-Mount (Cat#13800, LERNER labradorites) containing 5 μg/mL DAPI. Images were acquired with a 63× objective lens at 0.24 μm z-steps for a total of 17 slices using Axio Observer Z1 microscope equipped with Axiocam 506 mono camera (Carl Zeiss, Oberkochen, Germany). Acquired z-stacks were deconvolved using the constrained iterative algorithm in ZEN software (Carl Zeiss).

### Analysis of MAD1 localization

To quantify nucleoplasmic MAD1, a single cell was cropped using the same regions of interest (ROIs) from all channels (green, MAD1; red, POM121; blue, DAPI), and MAD1 signals outside the nuclear regions of interest (ROIs) were cleared. Using these cropped images, regions of MAD1 and POM121 signals were extracted based on fluorescence intensity thresholds and binarized. Colocalization was defined as the direct pixel overlap of MAD1- and POM121-positive pixels within the same ROI. MAD1-positive pixels colocalized with POM121-positive pixels were defined as NE-associated MAD1, whereas MAD1-positive pixels excluding POM121-positive pixels were defined as free MAD1.

Analysis of MAD1 localization was performed using ImageJ. MIP images were generated from the central five z-slices. Individual nuclei were cropped as square regions (200 × 200 to 250 × 250 pixels; 6.9383 µm/pixel) from all channels (green, MAD1; red, POM121; blue, DAPI). To quantify MAD1 localization during prophase, nuclear regions of interest (ROIs) were determined based on DAPI signals and expanded by 5 pixels to include perinuclear areas. Signals outside the expanded ROIs in MAD1 and POM121 channels were cleared. Prophase cells were identified based on chromosome morphology, showing condensed and heterogeneously stained DAPI signals with partial discontinuity at the nuclear boundary, as previously described (Kireeva et al., 2004). Binary masks of MAD1 and POM121 were generated using the Max Entropy automatic thresholding algorithm. The resulting binary masks were visually inspected and manually adjusted when necessary to correct obvious segmentation errors. Colocalization was defined as the direct pixel overlap of MAD1- and POM121-positive pixels within the same ROI. MAD1-positive pixels that colocalized with POM121-positive pixels were defined as NE-associated MAD1, whereas MAD1-positive pixels excluding POM121-positive pixels were defined as free MAD1. Release of MAD1 from the NE was quantified as the percentage of free MAD1 area relative to the total MAD1 area using the following formula: Occupancy (%) = (free MAD1 area) / (total MAD1) ×100.

### Analysis of IRE1 Activation

U2OS cells expressing IRE1-mNG under the control of a tetracycline-inducible promoter were seeded on poly-L-lysine–coated coverslips in 24-well plates. IRE1-mNG expression was induced with 2 µg/mL doxycycline for 48 hours (with medium replaced every 24 hours), after which cells were treated with 10 µg/mL Tm or 0.2 µM MG132 for the indicated times. Cells were rinsed with pre-warmed Hanks’ balanced salt solution (HBSS; Cat# 14025092; Gibco), and incubated at 37 °C for 30 minutes with 1 µM ER-Tracker Red (Cat# E34250; Invitrogen) in HBSS. Cells were then fixed with pre-warmed 4% PFA for 2 minutes at 37 °C and mounted on glass slides with 5 μg/mL DAPI.

IRE1 clustering was quantified from MIP images. The number of IRE1-mNG foci per cell was counted manually, and cells containing 10 or more foci were classified as IRE1-activated cells. At least 62 cells per condition were analyzed in three independent experiments.

### Western Blotting

For detection of ATF6, cells were lysed in SDS sample buffer (50 mM Tris-HCl, pH 6.8, 2% Sodium dodecyl sulfate (SDS), 10% glycerol, 10 μM MG132, and cOmplete Mini protease inhibitor cocktail [Cat# 11836170001; Roche, Basel, Switzerland]), boiled at 95°C for 5 minutes, and vortexed every 2 minutes. For other proteins, cells were lysed in lysis buffer (20 mM Tris-HCl, pH 7.5, 137 mM NaCl, 1% Triton X-100, 0.5% sodium deoxycholate, 0.1% SDS, 10% glycerol, 2 mM EDTA, 50 mM β-glycerophosphate, 1 mM sodium orthovanadate, 10 mM NaF, 1 mM dithiothreitol, 1 mM Phenylmethylsulfonyl fluoride, and 10 μg/mL leupeptin). Protein concentrations were determined using the Pierce BCA assay kit (Cat# 23225; Thermo Fisher Scientific). Equal amounts of total protein (30–50 μg) were separated by SDS-Polyacrylamide Gel Electrophoresis (SDS-PAGE) and transferred onto a polyvinylidene fluoride (PVDF) membrane (Cat# 88518; Thermo Fisher Scientific). Membranes were blocked with 5% nonfat milk in TBS-T (20 mM Tris–HCl, 150 mM NaCl, and 0.1% Tween-20) for 1 hour at RT, followed by incubation with primary antibodies and subsequently with HRP-conjugated secondary antibodies. Signals were detected using the SuperSignal West Pico PLUS Chemiluminescent Substrate (Cat# 34577; Thermo Fisher Scientific) and imaged with a ChemiDoc XRS+ system (Bio-Rad, Hercules, CA, USA). The following primary antibodies were used: anti-phospho-eIF2a (Ser51) (1:1,000; Cat# 3398; Cell Signaling Technology, Danvers, MA, USA), anti-eIF2α (1:1,000; Cat# sc-11386; SantaCruz), anti-ATF6 (1:2,000; Cat# 24169-1-AP; ProteinTech), and anti-GAPDH (1:1,000; Cat# 5174; Cell Signaling Technology). The following secondary antibodies were used: Goat anti-Rabbit IgG (H+L) Secondary Antibody, HRP (1:5,000; Cat# #32460; Invitrogen) and Goat Anti-Mouse IgG (H+L)-HRP Conjugate (1:5,000; Cat# 170-6516; Bio-Rad).

### Statistical Analysis

Statistical analyses were performed using Student’s t-test or analysis of variance (ANOVA), followed by Sidak’s multiple comparisons test when applicable. For cumulative distribution plots, statistical significance was assessed using the chi-square (χ²) test. A *p*-value < 0.05 was regarded as statistically significant.

